# Transit-amplifying progenitor with maturation-dependent behavior contributes to epidermal renewal

**DOI:** 10.1101/2022.06.12.495812

**Authors:** Sangeeta Ghuwalewala, Kevin Jiang, Sara Ragi, David Shalloway, Tudorita Tumbar

**Affiliations:** Department of Molecular Biology and Genetics, Cornell University, Ithaca, NY, USA

## Abstract

Transit-amplifying progenitor populations with phased behavior have long been postulated as essential to epidermal renewal, but not experimentally observed *in vivo*. Here we identify a population with bi-phasic behavior using CreER genetic cell-marking in mice for long-term lineage tracing and clonal analysis. Nascent, highly expressing Aspm cells undergo an amplification-phase followed by a timed transition into an extinction-phase, with near complete loss of descending cells from skin. Generalized birth-death modeling of Aspm-CreER and a Dlx1-CreER population that behaves like a stem cell demonstrates neutral competition for both populations, but neutral drift only for the stem cells. This work identifies a long-missing class of non-self-renewing epidermal progenitors with bi-phasic behavior that appears time-dependent as the lineage matures, indicative of a transit-amplifying cell. This has broad implications for understanding cell fate decisions and tissue renewal mechanisms.

**One-sentence summary:** We identify a long-missing class of non-self-renewing epidermal progenitors with bi-phasic and maturation-dependent behavior in vivo.

## Introduction

Adult epidermal renewal is essential for the skin barrier function and occurs from proliferative cells located in the basal layer (BL) that differentiate upwards into the supra-basal layers (sBL) until they are shed out from the skin surface (*1*). Despites decades of work, the lineage organization of stem cells (SCs) and progenitor cells in this tissue is not yet fully established. Some lineage organization models based on live imaging and genetic marking driven from ubiquitous promoters implicate a single equipotent progenitor, with balanced and stochastic choices to self-renew and differentiate known as neutral drift (*1–3*) (*4–6*). Other studies support a two-population hierarchical model with SCs (marked by K14-CreER) generating a committed progenitor (marked by Inv-CreER); both populations undergo stochastic choices of self-renewal vs differentiation that are continuously imbalanced towards self-renewal for the SCs and towards differentiation for progenitors (*7–9*) (Fig. S1A,B). Most recently, another highly committed and very short-lived (~1-week) population of BL progenitors that express the differentiation marker K10 already and divide only 1-2 times before exit into sBL has been described by both lineage tracing and life imaging (*10*). Importantly, all of these stem/progenitor cells described to date in the basal layer have displayed constant or continuous growth properties. Specifically, cell fate choices identified by lineage tracing are either: i) constantly balanced between renewal and differentiation to maintain the pool of long-term self-renewing (SR) progenitors, referred to as stem cells (SCs) or ii) constantly unbalanced towards differentiation for non-self-renewing (NSR) progenitors.

The <u>constant</u> growth properties of SR and NSR progenitors identified to date *in vivo* contrast with the classical behavior of cultured untransformed primary cells, which undergo phased or maturation-dependent behavior that changes over time (*11*). Many primary cultured cells undergo bi-phasic behavior. First after plating, as they are young, they undergo a growth or *amplification phase* when they divide and increase their numbers. Then as they ‘mature’ these cells reach a proliferation limit (i.e., the Hayflick limit) when they switch into an *extinction phase* and eventually die out (*11*). In addition, primary cultured human epidermal cells have distinct proliferation limits (e.g. form holoclones, meroclones, and paraclones) most recently associated with specific transcriptomic profiles (*12*), suggesting distinct potentials as long-lived SCs vs short-lived progenitor cells *in vivo*. Importantly, maturation-dependent bi-phasic behavior of epidermal progenitors (Fig. S1C)—transit-amplifying (TA) cells—has not yet been demonstrated *in vivo*. Decades-old predictions from classical tissue kinetics studies suggested a hierarchical model, with slow-cycling SCs generating frequently dividing TA cells that then terminally differentiate (TD), with important implications for tissue aging and cancer (*13–15*).

Here, we experimentally identify a type of epidermal NSR TA progenitor in mouse tail skin *in vivo* marked by the Aspm-CreER genetic driver. This population undergoes a bi-phasic behavior consistent with an amplification followed by an extinction phase. We analyze this behavior implementing a generalization of ‘birth-death modeling’ (*16*) to lineage tracing data and show that it preserves neutral competition without neutral drift. This behavior directly contrasts to that of another, less proliferative epidermal population marked by Dlx1-CreER, which undergoes neutral drift. This work bridges old tissue kinetics and classical SC-TA-TD theory (*13–15*) with modern clonal evolution and live imaging analysis (*1–3, 5, 7, 10, 17*) and uncovers a new population of epidermal progenitor with bi-phasic behavior that changes as this lineage matures over time.

## Results

### Basal layer cells with high Aspm expression are predicted short-term epidermal progenitors

Anticipating that TA cells may proliferate faster than SCs (*13–15*), we previously employed H2B-GFP pulse-chase transgenic mice (*18*) to isolate basal layer (BL) cellular subsets with distinct proliferative properties. We found that Aspm and Dlx1 mRNAs were upregulated in faster and slower dividing BL cells, respectively (*19*). The Dlx1-CreER genetic driver marked infrequently dividing long-lived epidermal progenitors (*19*), suggesting that the Dlx1-CreER cells contain SCs; however, clonal analysis was not performed. Here we employ these two markers in clonal lineage tracing experiments to examine their behavior as putative TA cells and SCs.

Single cell (sc) transcriptomics using 10x Illumina RNA-sequencing of our Sca1^+^/α6-integrin BL cells sorted from mouse tail skin (*20*) revealed that the Aspm and Dlx1 mRNAs are expressed in different subsets of BL cells, and are both largely distinct from differentiating K10+/K14+ BL cells (Fig. 1A-D). Moreover, Aspm+ (but not Dlx1+) cells were highly enriched in the S/G2/M BL clusters and were >90% Ki67+, demonstrating their strong proliferative status (Fig. 1A-E). We recently reported Aspm expressed at the protein level in a fraction on BL cells in mouse tail and back skin and in human skin (*20*). Lineage trajectory prediction using Monocle 2 of the sc transcriptomics data revealed the putative ‘ground state’, which was least enriched in differentiation marker K10 and was interestingly enriched in Aspm+ cells (Fig. 1F). This is unlikely to be an attribute of Aspm expression in a subset of Ki67^+^ proliferative cells alone, since Ki67^+^ cells formed a highly branched, split-lineage tree that was not assigned a ground state in related hair follicle SC populations (*21*). These data suggest that high Aspm expression may be characteristic of an important highly proliferative short-term progenitor, a working-horse of epidermal renewal.

**Figure 1.**
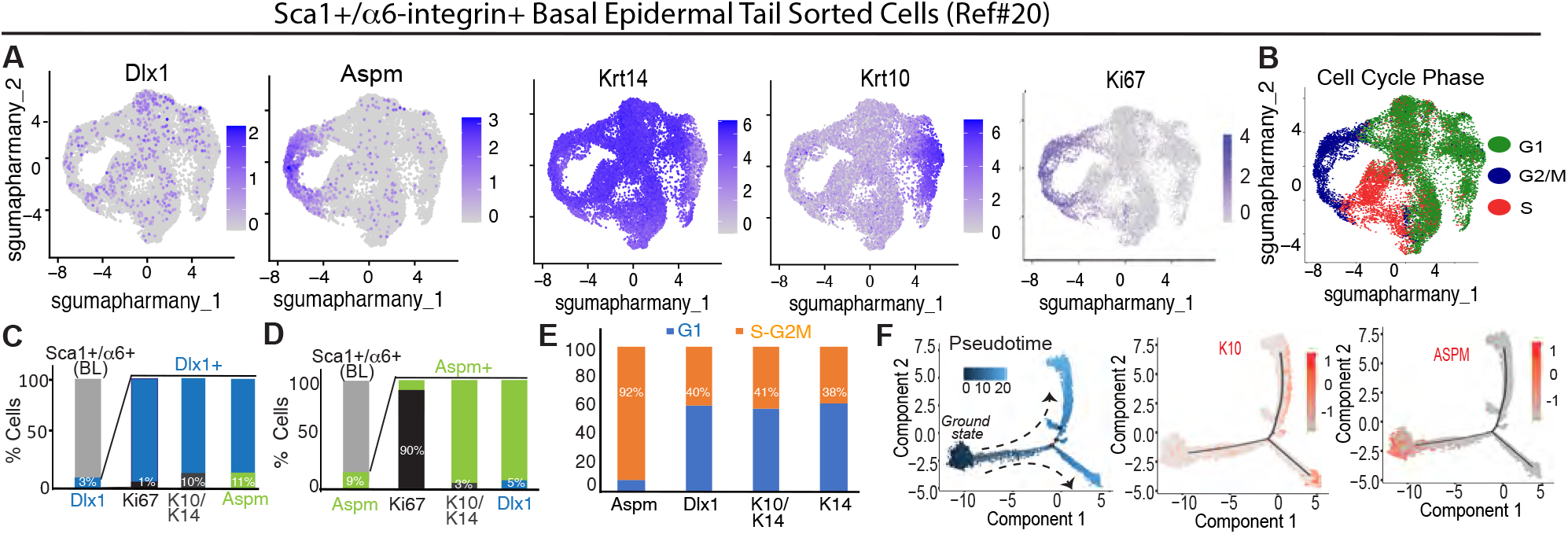
Aspm and Dlx high-expressing basal cells mark IFE populations with distinct properties. (A) scRNAseq feature plots of Sca1+/a6-integrin+ basal layer (BL) cells sorted from mouse tail skin. Shades of blue show expression levels. (B) Regression analysis of scRNAseq data shows distribution of cell cycle phases in BL cells. (C,D). Quantification of scRNAseq BL data in (A) demonstrate little overlap of Dlx1+/Aspm+/K10+ cells and high overlap for Aspm/Ki67+ cells. (E) Fraction of distinct basal cell populations in cell cycle phases demonstrate 92% Aspm+ cells are actively cycling (in S/G2/M). (F) Comparative lineage trajectory with Pseudotime analysis using Monocle 2 predicting progenitor ground state with K10 and Aspm expression feature plots. Dlx1 cannot be featured on Monocle 2 graphs due to low expression levels.

### Dlx1-CreER marks stem cells and Aspm-CreER marks bi-phasic progenitors with changing behavior during lineage maturation

To examine the behavior of Aspm- and Dlx1-expressing epidermal populations during homeostatic epidermal renewal, we performed genetic clonal labeling and lineage tracing up to one year (Fig. 2A). We used Dlx1-CreER as previously reported (*19*) and Aspm-CreER transgenic mice (*22*) crossed with the Rosa-26-tdTomato reporter. Low tamoxifen (TM) doses ensured targeting rare basal cells for clonal analysis. When the Aspm-CreER mice were induced with high tamoxifen (TM) dose and 5-daily injections followed by long-term chases the tail epidermis was highly efficiently labeled (Fig. S2).

**Figure 2.**
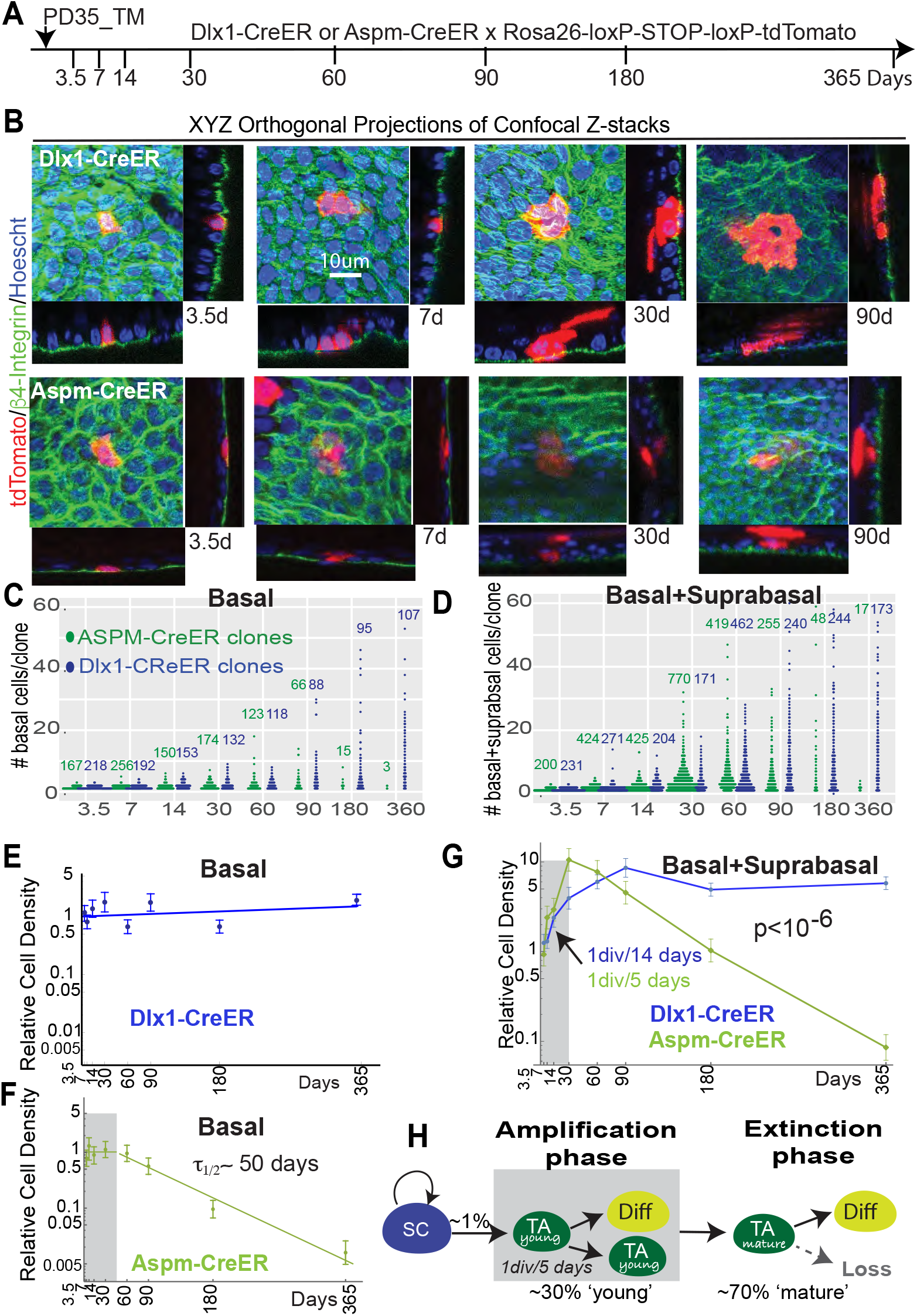
Genetic lineage tracing reveals a stem cell and a maturation-dependent progenitor with biphasic behavior. (A) Low dose (clonal) tamoxifen (TM) induction with chase times indicated. (B). Examples of clones from the two lineages show tdTomato marked cells in the basal (β4-integrin^+^) vs suprabasal locations. (C,D). Summary of tdTomato^+^ cell counts shown as beeswarm plots, collected from comparable tail skin areas (Table S1). Number at top indicate total number of clones counted. (E,F) Relative BL cell densities as a function of chase-time with best log-linear fits to the data. For Aspm-CreER, amplification-phase (gray box) and extinction-phase are fit separately. (G) Relative total cell densities. (H) Inferences from the Aspm-CreER density analysis.

For clonal analysis, whole-mount tail skin collected from the two mouse lines at timepoints indicated were stained with BL marker β4-integrin and imaged by xyz-automated high resolution confocal microscopy (Fig. 2B and Table S1). The tdTomato^+^ clones and cells in both BL and (differentiated) supra-BL (sBL) layers from different mice were quantified (Fig. 2C,D and confirmed separately with low resolution counts in Fig. S3). Over 2,500 clones in the Aspm-CreER and 1,995 in the Dlx1-CreER populations were assayed over the 8 experimental timepoints (Fig. 2C,D and Table S1). The clone size distribution did not reveal even-number bias, which was previously reported in mouse back skin and was attributed to stem-progenitor paired behavior (*9*). Our results that lack even-number bias were in line with other previously reported mouse tail clonal data (*7, 8*).

The cell densities (normalized cells/image) of the two populations evolved during the year in notably different ways, indicative of distinct progenitor cell types (Fig. 2 E-G). Specifically, Dlx1-CreER-marked cells largely maintained BL density over one year, while contributing to differentiated sBL cells, indicative of a self-renewing SC population (Figs. 2E,G and S1A). In contrast, Aspm-CreER-marked cells contributed robustly to sBL while maintaining BL density for ~ 30 days in an initial phase. We refer to cells in this initial phase as ‘young’, to illustrate the early stages of the chase and their robust proliferative behavior. Around 30-60 days of chase the Aspm-CreER-marked cell numbers began an exponential decline, entering a late ‘extinction-phase’, with a half-life of ~50 days and near complete long-term loss from skin by one year (Figs. 2F,G and S1C). We refer to cells in this second phase as ‘mature’, to illustrate both the late chase times as well as their extinction-prone behavior. The rate-of-increase of BL+sBL density in the early phase (i.e., up to 30 days) implied that Aspm-CreER cells divided once every ~5 days, which was ~3x faster than Dlx1-CreER (Figs. 2G and S4, Table S2), as expected from our mRNA expression (Fig. 1) and our previous studies that identified Dlx1 and Aspm as putative markers (*19*). The sBL labelled cell densities at late timepoints depend on the unknown dynamics of cell-shedding from outermost skin layer, and so could not be used for total rate calculations during the interval past 30 days. The behavior of the two lineage-traced populations did not differ in spatially distinct domains of the tail skin, known as scales and inter-scales (Fig. S5).

Assuming homeostatic maintenance, the BL density curves (Fig. 2E,F) imply that ~1%/day of the homeostatic population is replenished by input from a SC or other epidermal population (see Supplementary Theory, ST). Moreover, we calculate that approximately 30% of BL cells in the homeostatic population are ‘young’, undergoing the ‘amplification-phase’ (see Figs. 3 and 4), and 70% are ‘mature’ and undergoing extinction (Fig. 2H and ST, Sec. II).

**Fig. 3.**
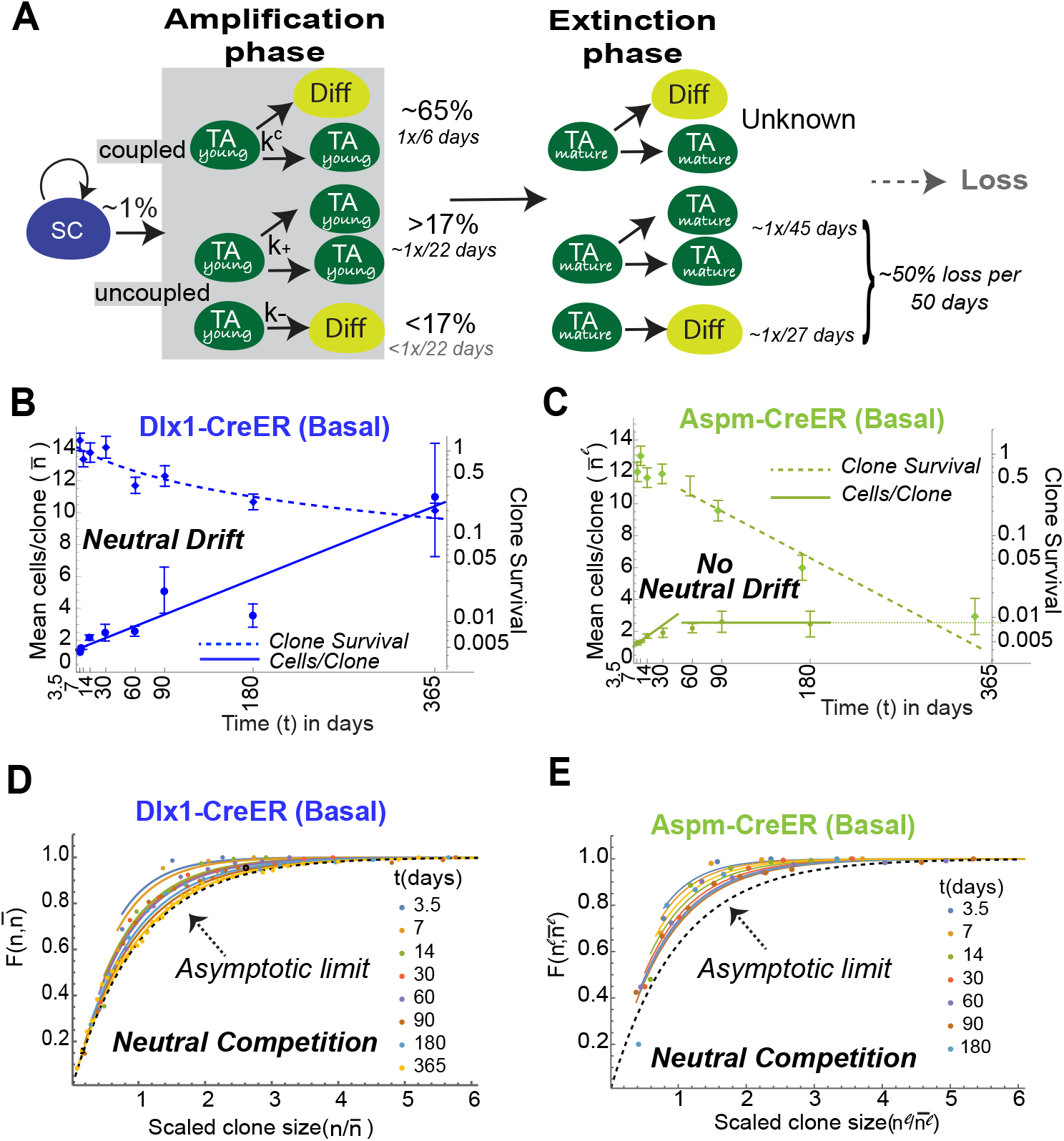
Generalized birth-death modeling demonstrates neutral competition without neutral drift of maturation-dependent progenitor. (A). Division and differentiation/export rates calculated for Aspm-CreER. Rates of coupled division/export (*k^c^*), uncoupled division (*k*_+_), and uncoupled export (*k*_−_) in amplification phase were calculated using the cell density, clone-size, and clone survival lineage tracing data. *k*_+_ > *k*_−_in amplification phase and *k*_−_<*k*_+_ in extinction phase (Table S2 and ST, Sec. II.C for details). TA, transit amplifying progenitor. (B) Hyperbolic decrease in clone survival and linear increase in mean clone-size of Dlx1-CreER consistent with neutral drift. (C) Exponential decrease in clone survival and finite mean clone-size maximum of Aspm-CreER not consistent with neutral drift. (D,E) Scaled cumulative plots of clone-size *n* at the chase timepoints. 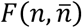 is the fraction of clones with clone-size ≤ *n*, where 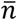 is the mean clone-size at that timepoint. Superscripts ^*1*^ indicate that the observed Aspm-CreER values depend on the variation of its labeling-efficiency with progenitor cell maturity (time after its ‘birth’ in BL). The solid lines are the neutral competition predictions and provide excellent fits to both the Dlx1-CreER and Aspm-CreER clones. The asymptotic ‘scaling’ limit at large chase-time predicted for clones undergoing neutral drift (dashed line) is attained by Dlx1-CreER, but not Aspm-CreER. The individual plots are displayed in Fig. S6.

**Fig. 4.**
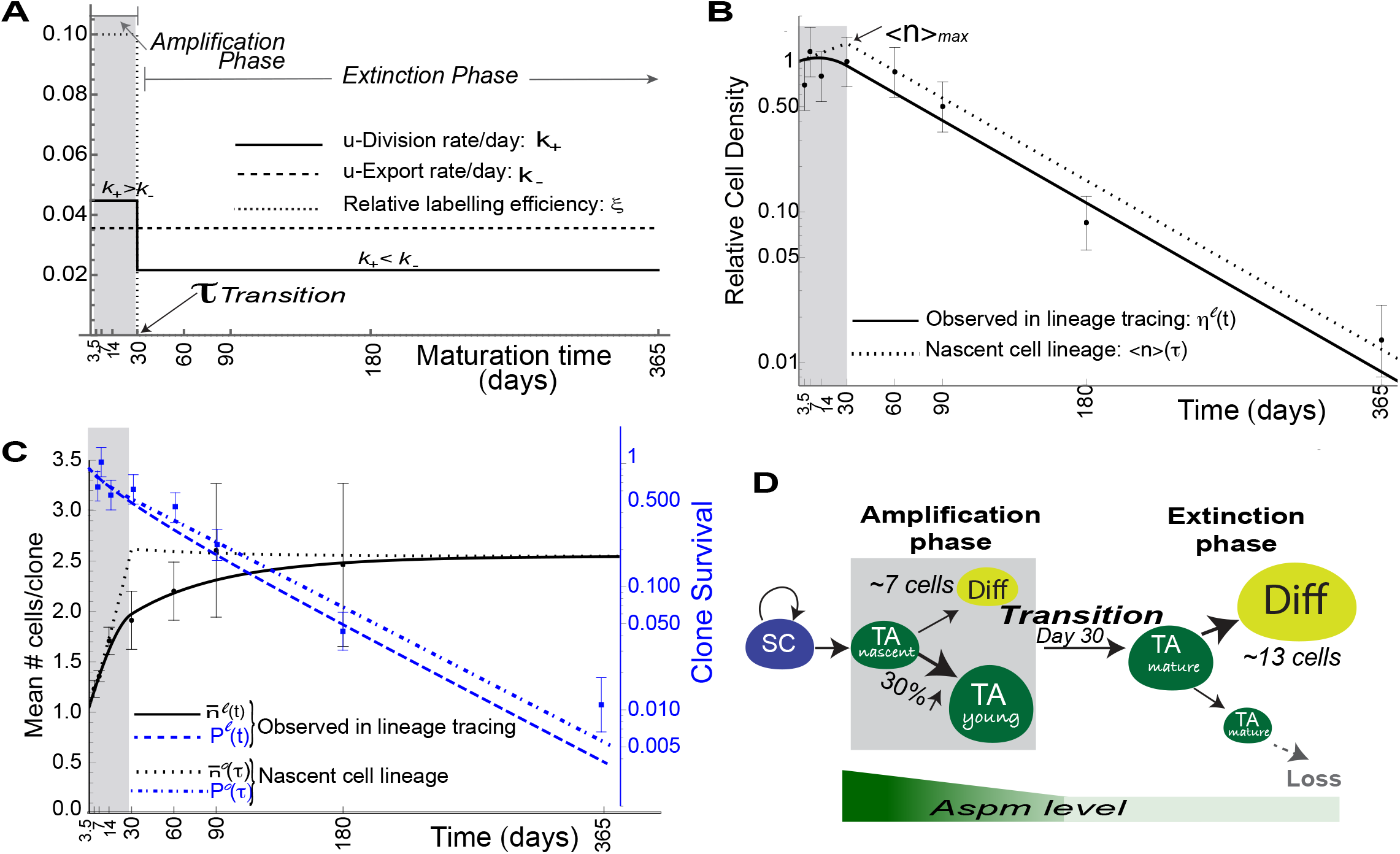
‘Timed-transition’ model for Aspm-CreER transit amplifying (TA) progenitor behavior. (A) Maturation-dependence of parameters used in the model. (B,C,D) Markers and solid and dashed lines are the observed, labeled data with model predictions over chase-time *t*. Dotted lines are the model predictions for 〈*n*〉(τ), mean number of cells in a lineage; *P*^0^(τ), survival probability of a nascent cell’s lineage; 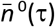, the mean size of surviving clones. τ is lineage maturity. (D) The mean size of a nascent cell’s lineage increases by 30% and ~7 cells terminally differentiated (Diff) are exported to sBL during amplification-phase. Assuming the ratio of coupled to uncoupled division remains the same, an additional ~17 terminally differentiate (Diff) cells are made in extinction-phase. See ST and Table S3 for mathematical notation and calculation. Superscripts^*ℓ*^ denote dependence of observed labelingefficiency on Aspm-CreER maturity. See ST, Sec. II.D and Table S3 for calculations.

We conclude that Dlx1-CreER acts as a bona fide long-term self-renewing (SR), slow-cycling SC, while Aspm-CreER marks a more rapidly dividing non-self-renewing (NSR) progenitor. The latter displays a biphasic behavior that changes as the lineage matures over time, suggestive of a distinct transit-amplifying (TA) cell population (Fig. 2H). The findings remind of early lineage-dynamics based on tissue kinetics that implicate both SCs and TA cells with maturation-dependent behavior(*13–15*), and prompt us to further inquire into the nature of cell fate choices in the two epidermal populations.

### Neutral drift clonal dynamics for Dlx1-CreER SCs, but only neutral competition for Aspm-CreER TAs

While historical tissue kinetics data suggested non-random, linked SC®TA® TD cell fates in a hierarchical organization (*13–15*), modern clonal analyses and live imaging data support stochastic cell fate choices by equipotent cells (i.e., neutral competition) between division and terminal differentiation (e.g. differentiation with export into sBL) (*1–3, 5*). As previously described for modeling of clonal analysis cell division and loss from the basal into the supra-basal layer (e.g. export) (*2, 4, 8*) can be either ‘coupled’ and/or ‘uncoupled’ events that contribute to overall tissue growth (Fig. 3A, grey panel). ‘Coupled’ division and export (often referred to as ‘asymmetric cell fates’) are temporally correlated events within a clone on a time-scale commensurate with their rates and could result from asymmetric cell divisions (*23*) or symmetric divisions linked with cell delamination from the BL (*5*). For the most part, only the uncoupled events affect BL cell density, clone survival, and clone size and, unlike the coupled events, can have unequal rates resulting in non-self-renewal (NSR). (For notational simplicity, we group the possible event where both daughter cells from a division are immediately exported with the ‘uncoupled’ events.) In a self-renewing (SR) progenitor population (e.g. SCs), the rate of cell division equals that of export (e.g. are ‘balanced’) and, in lineage tracing, the number of observed (or labelled) clones decreases while the sizes of the observed surviving clones increase so as to maintain overall cell number. This is known as ‘neutral drift’ and is predicted as consequence of stochastic critical birthdeath processes (*1, 24*). Consistent with neutral drift, the Dlx1-CreER mean BL clone-size increased linearly with chase-time, while clone survival decreased hyperbolically—the product giving constant BL cell-density (Fig. 3B). In contrast, after increasing in amplification-phase, the Aspm-CreER BL mean clone-size stabilized and clone survival decreased exponentially in extinction-phase, violating neutral drift (Fig. 3C).

The ‘critical birth-death’ theory that was previously used to predict neutral drift is not applicable when the division and export rates are imbalanced and change with maturation, as with Aspm-CreER. Moreover, if the experimental labeling-efficiency varies in TA cells as a function of their ‘maturity’ (for instance, CreER preferentially marks ‘young’ vs ‘mature’ cells or vice-versa), this can complicate the relationship between observations and the underlying biological properties. To meet this challenge, we developed a ‘generalized birth-death theory’ that accommodates NSR, maturation-dependent rates, and maturation-dependent labeling efficiency (see ST).

Using this we showed that, while Aspm-CreER clones do not display neutral drift, clonal evolution is still consistent with the weaker condition of cell-autonomous neutral competition —i.e. stochastic division and loss from basal layer (e.g.export). This predicts that the fraction of BL clones smaller than size *n*—i.e., the cumulative distribution—is

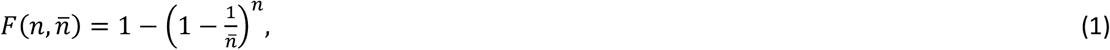

(where 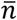 is the mean clone-size at the time of measurement) when rates are constant, and is a good approximation even when they are bi-phasic or ‘maturation-dependent’ (see ST, Sec. I.B.1,2). If 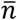 gets large, this converges to the asymptotic ‘scaling’ limit 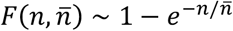, which is a function only of 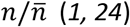, and is best examined in scaled plots over this ratio (Figs. 3D,E and S6A,B; ST, Sec. I.B.1). Both the Dlx1-CreER and Aspm-CreER distributions agreed at all timepoints with this prediction, consistent with neutral competition. The Dlx1-CreER distributions converged at large chase-times to the scaling limit (dashed line) as expected for neutral drift, but the Aspm-CreER distributions did not because their mean clone-sizes remained small.

This analysis demonstrates that not only SR SCs (*1, 5*), but also maturation-dependent TA cells with bi-phasic behavior can undergo independent, stochastic division vs TD (i.e., differentiation/export) choices during both the amplification and the extinction phases. This contradicts the historical non-stochastic SC®TA® TD cell hierarchical model predicted by tissue kinetics studies (*13–15*) and extends current SC theory (*1, 5*) to TA cells, which display neutral competition without neutral drift.

### Cell division and differentiation/export of Dlx1-CreER and Aspm-CreER subpopulations

The constancy of the SR Dlx1-CreER BL cell density implies that the uncoupled (u)-division and u-export rates—*k*_+_ and *k*_−_, respectively—are equal and can be determined using the standard critical birth-death theory from linear and hyperbolic fits to the Dlx1-CreER BL mean clone-size and clone survival, respectively (Fig. 3 B). This gives one u-division and one u-export every ~45 days (, Table S2, ST, Sec. II.B). This represents a fraction of the total initial divisions (coupled+uncoupled) that occur once every 15 days (Fig. 2G and S4).

Because of the varying, imbalanced proliferation and export rates, analyzing the Aspm-CreER data required the generalized theory (ST). The initial rate-of-increase of mean clone-size (Fig. 3C) implied that these cells had one u-division every ~22 days during amplification phase. This is ~4× slower than the amplification-phase total division rate (Figs. 2G,H and Table S2), implying that coupled (= total-uncoupled) divisions occur every ~6 days and comprise ~70–85% of all the Aspm-CreER divisions. The relative frequencies of all the coupled and uncoupled events in amplification-phase are summarized in Fig. 3A. Using the asymptotic value of the mean clone-size and the rate of exponential-decrease, we calculated that there was one u-export every ~28 days but only one u-division every ~45 days in extinction-phase (ST). This creates an unbalance towards differentiation that explains the loss of marked cells in the long-term (Fig. 3A). Accounting for effects of the ~2× decrease in u-division rate with maturation, the theory predicts that there is an increase in BL density during amplification-phase that is masked by the experimental error (ST and Fig. 3C).

### A timed-transition model predicts TA expansion of BL cells during amplification phase

In a progenitor population with bi-phasic or ‘maturation-dependent’ behavior the clonal evolution observed in lineage tracing depends on which cells of the lineage are targeted by the initial labeling. For instance, if the CreER is activated equally in cells of all lineage ages (e.g. young and mature), then a continuous decay in the number of initially labeled cells would be observed, until all young and mature cells are lost. This type of clonal behavior was reported for the Inv-CreER tail population (*8*).

Alternatively, if only the mature cells are labeled, a quick loss from the skin of the labeled population would be observed, as was the case for the K10+/K14+ marked basal population (*10*). In our data, the observed bi-phasic behavior of the Aspm-CreER marked cells can only be explained if ‘young’ TA cells are preferentially labeled (see ST for theoretical analysis). This labeling bias can have multiple experimental explanations, of which the simplest is that ‘young’ cells express the highest Aspm levels and therefore the CreER is preferentially induced in these cells by the low TM injections used for clonal analysis.

It is interesting to consider a specific model that is consistent with the conclusions drawn above, although other models cannot be excluded. To avoid overfitting, we made several mathematical approximations: i) the transition from young to mature occurs exactly at 30 days; ii) the u-export rate is constant, while the u-division rate decreases at 30 days; and iii) the labeling preference for young cell is absolute (Fig. 4A). After fitting three adjustable parameters (Table S3), the model matches the observed BL data and clone-size distributions over the entire chase period (Figs. 4B–D and S6C). This ‘timed-transition model’ predicts that the mean number of BL cells in a nascent cell’s lineage expands ~30% and that ~7 cells are exported to sBL during 30-days amplification phase (Fig. 4D). If the ratio of coupled-to-uncoupled divisions remains constant, then ~13 more differentiated cells are exported during the long extinction-phase (Fig. 4D).

In summary, using Aspm as a novel epidermal progenitor marker for clonal analysis together with a new application of generalized birth-death modeling to lineage tracing, we identify an epidermal transit amplifying (TA) cell in vivo that has bi-phasic behavior, which changes as the lineage matures over time. We find that our nascent TA cells have an initial amplification-phase and then, ~30 days later, transit a biological turning point from a cell state more prone to proliferation to one more prone to differentiation. This discontinuous timed-transition is not consistent with a recently proposed model of gradual differentiation for BL cells (*10*) (*25*). With that said, previously proposed gradual differentiation models referred to the last stage of BL differentiation where already committed K10+ basal cells exit into the sBL (*10, 25*); the gradualistic models do not account for less committed K10-progenitors, such as our TA cells, and switches of cell states within the BL, as is the case here.

Our data and modeling suggest that during their two phases, TA cells undergo stochastic fate choices between cell division and differentiation/export into the sBL; these appear imbalanced towards division in amplification-phase and towards differentiation/export in extinction-phase. The bi-phasic behavior of our NSR TA population distinguishes it from the previously described NSR Inv-CreER committed progenitor (CP) population (*7, 8*), which is constantly imbalanced towards differentiation, thus displaying a continuous loss of labelled cells over time.

Using a generalized birth-death model implemented first time to clonal analysis, we described in depth the cell kinetics of proliferation and differentiation/export of our Aspm-CreER TA population. Our TA cells divide on average 1x/5 days, similarly with the bulk of BL cells (*13–15, 26*); this rate is 3x faster than our control SC population marked by Dlx1-CreER, in agreement with the classical slow-cycling SC-rapidly dividing TA cell model. We find that the Aspm-CreER TA population is much longer-lived (several months) than suggested for TA cells from early epidermal tissue kinetics studies (2-3 weeks) (*15, 26*) and from life-imaging studies (1-2 weeks) (*10*). This may explain why our population remained hidden in short-term live imaging studies (*5*), since Aspm-CreER TAs and SCs behave similarly over weeks-long time intervals. In fact, as predicted decades ago (*14, 15, 26*), it appears that multiple TA cell states with distinct proliferation and differential potential, intermediate between a SR SCs and terminally differentiated cell types exist in the BL.

To maintain homeostasis, half of the Aspm-CreER TA population is replaced every ~3 months from another epidermal progenitor population. We do not know which epidermal cells replenish the Aspm-CreER-marked cells, but any basal SC population such as Dlx1-CreER (this work) or K14-CreER (*8*) marked cells could be a cell source. Alternatively, the Aspm-CreER population may not be replenished in old age past 1-year old being present only in young skin, a possibility that should be investigated in the future. Regardless of these considerations, our results open up new avenues into investigating mechanisms that distinguish SR and NSR epidermal cell types *in vivo* and of timed-transitions from young to mature TA cell states. As inferred by early historical models (*11, 14*), these mechanisms have broad implications, not only in cell fate decisions and homeostatic renewal but also in tissue aging and cancer.

## Acknowledgments

We thank P.A. Schweitzer for generating 10x genomics scRNA-seq data; C. J. Bayles, R. M. Williams and J. M. DelaCruz for helping with confocal imaging (BRC Facility) and data processing; and the Cornell CARE staff for mouse husbandry. The Cornell Biotechnology Research Center and Imaging Facility as supported by NIH Grant 1S10RR025502-01. The research was supported by grants from the National Institute of Arthritis and Musculoskeletal and Skin Diseases (R01AR070157 and R01AR073806) to TT and the CVG scholar grant 2021 awarded to SG.

## Author contributions

S.G., D.S. and T.T. designed the experiments and interpreted the data. S.G., K.J. and S.R. performed experiments and clonal analysis. S.G, D.S. and T.T. made the figures and wrote the manuscript. S.G. and D.S. performed statistical analysis and generated graphs. D.S. adapted and applied generalized birth-death theory to lineage tracing.

## SUPPLEMENTARY FIGURE LEGEND

**Fig. S1.**
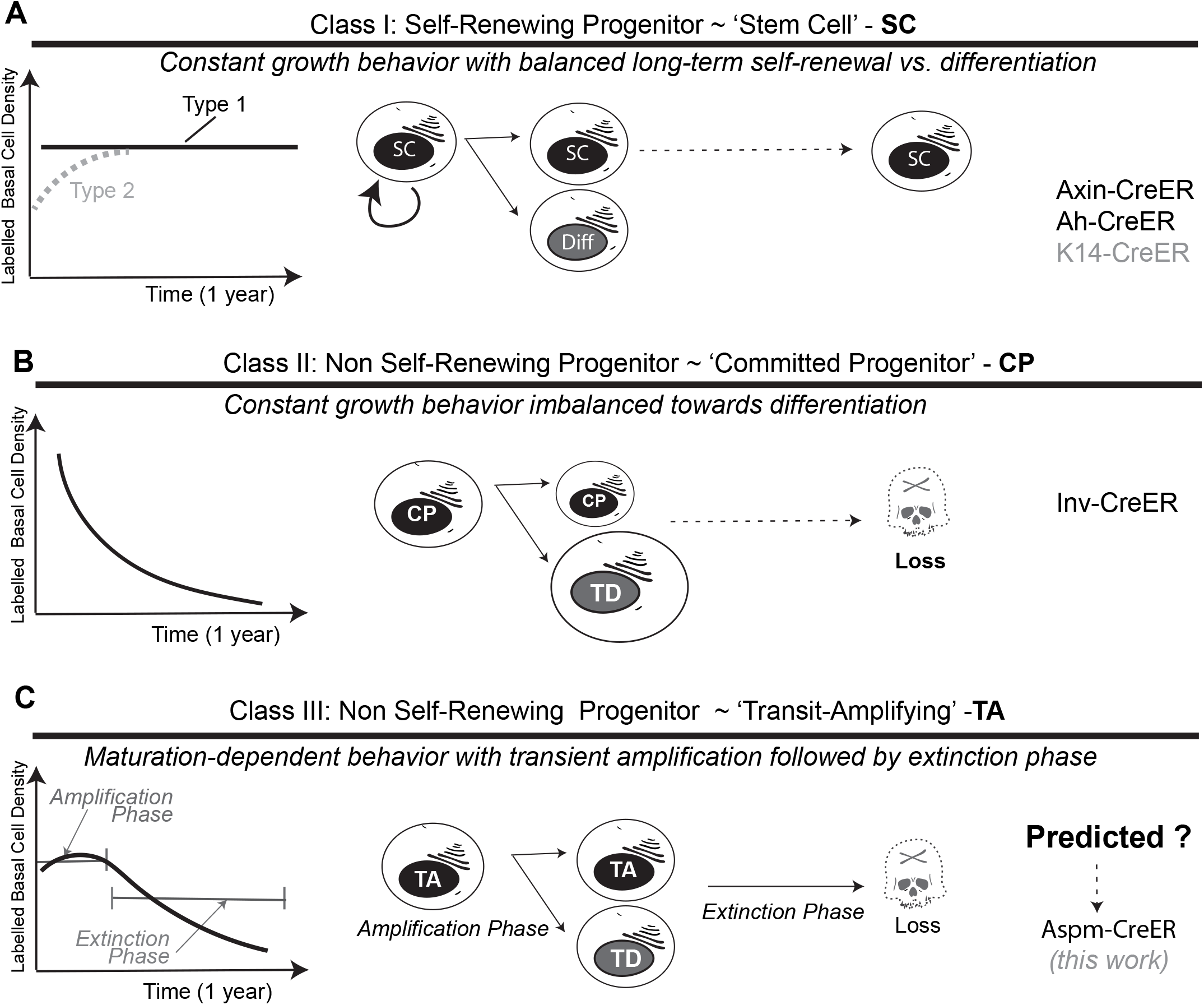
Classes of epidermal progenitors and their behavior in homeostasis. (A) A self-renewing (SR) stem cell (SC) population displays basal cell density detected in lineage tracing either as a constant (Type 1, Axin-CreER (*27*) and Ah-CreER (*2*)), or as an asymptotic increase to steadystate, if the labeled lineage accumulates descendants (Type 2, K14-CreER (*7, 8*)). (B) A non-self-renewing (NSR) ‘committed’ progenitor (CP) with a constant imbalance towards differentiation (e.g. Inv-CreER (*7, 8*)) displays an exponential decrease of basal cell density in lineage tracing. (C) A maturation-dependent bi-phasic behavior (e.g. a transit-amplifying (TA) progenitor) displays an amplification and an extinction phase, which are observable in lineage tracing if ‘young’ cells are preferentially labeled overed mature cells. Aspm-CreER described in this paper is the first epidermal progenitor of this type identified *in vivo*.

**Figure S2.**
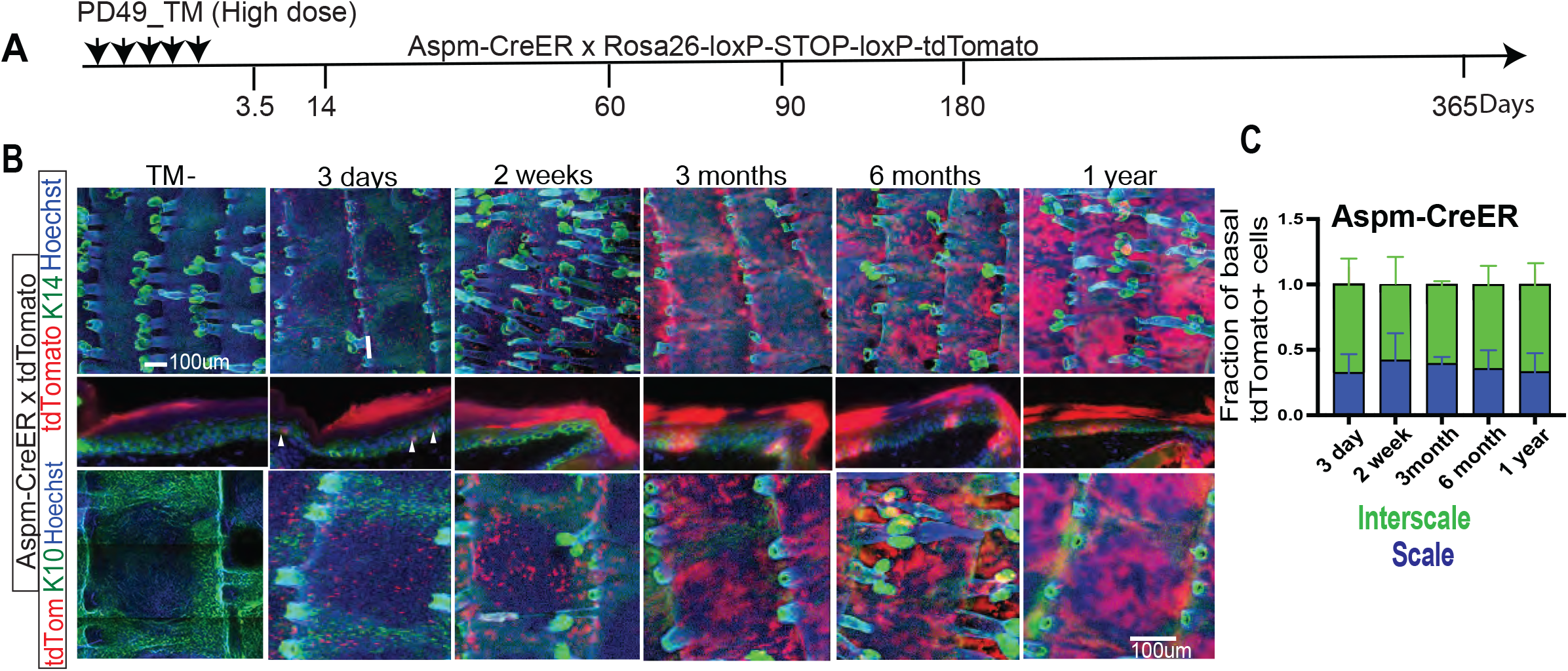
High dose tamoxifen induction of Aspm-CreER efficiently labels the epidermis in long-term lineage tracing. (A) Schematic of mouse tamoxifen (TM) induction and chase timeline with collected time points indicated. (B) Whole mount skin tissue IF stained with antibodies as indicated are visualized as maximal projections from confocal microscopy image stacks shown at time points and with magnification indicated. (C) Quantification of images like those in (B) from n=2 mice show the distribution of tdTomato labelled basal cells in both scale and inter-scale tail skin regions. Hoescht is DNA staining (blue). Scale bar, 100μm.

**Figure S3.**
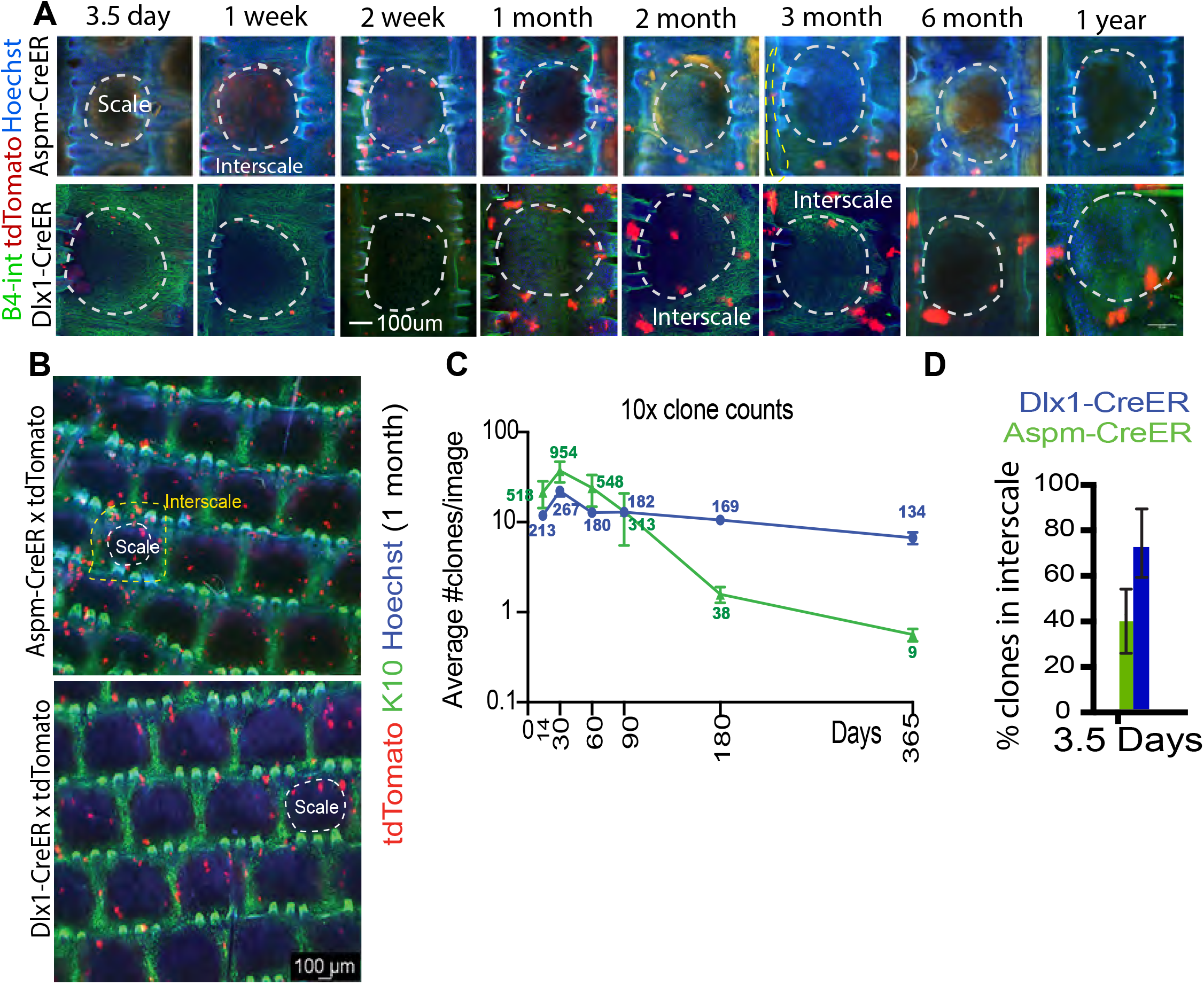
Long-term clonal lineage tracing in tail skin. (A) Confocal images show examples of clones at different time points indicated in one scale area with its associated interscale area (Scale bar, 100um). (B) Widefield fluorescence with 10x magnification objective shows tail whole-mount immunofluorescence stained for K10 marking the interscale from 1-month lineage-traced mice of genotype indicated. (C) Clone counts of 10x images at time points indicated. Total number of clones identified at each time point are shown. (n>2 mice and N>5 images per each time point). (D) Fraction of clones in scale vs interscale at PD3.5 extracted from the high-resolution 3D confocal data (Table S1). Note higher fraction of Dlx1-CreER clones in the interscale as previously reported (*19*). Blue is DNA DAPI staining.

**Fig. S4.**
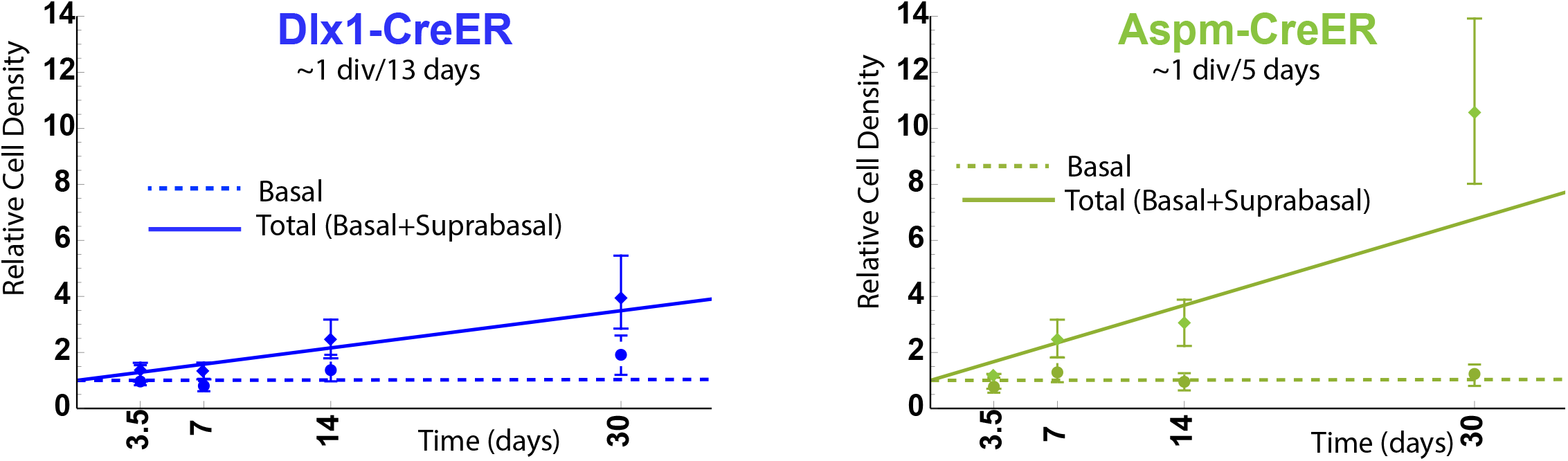
Total initial cell division rates in amplification-phase. The least-squares fits to the total cell densities for the first 30 days chase of basal and suprabasal were computed (solid) and compared with the BL fits from Fig. 2E,F (dashed) used to compute the total division rates shown at the top.

**Fig. S5.**
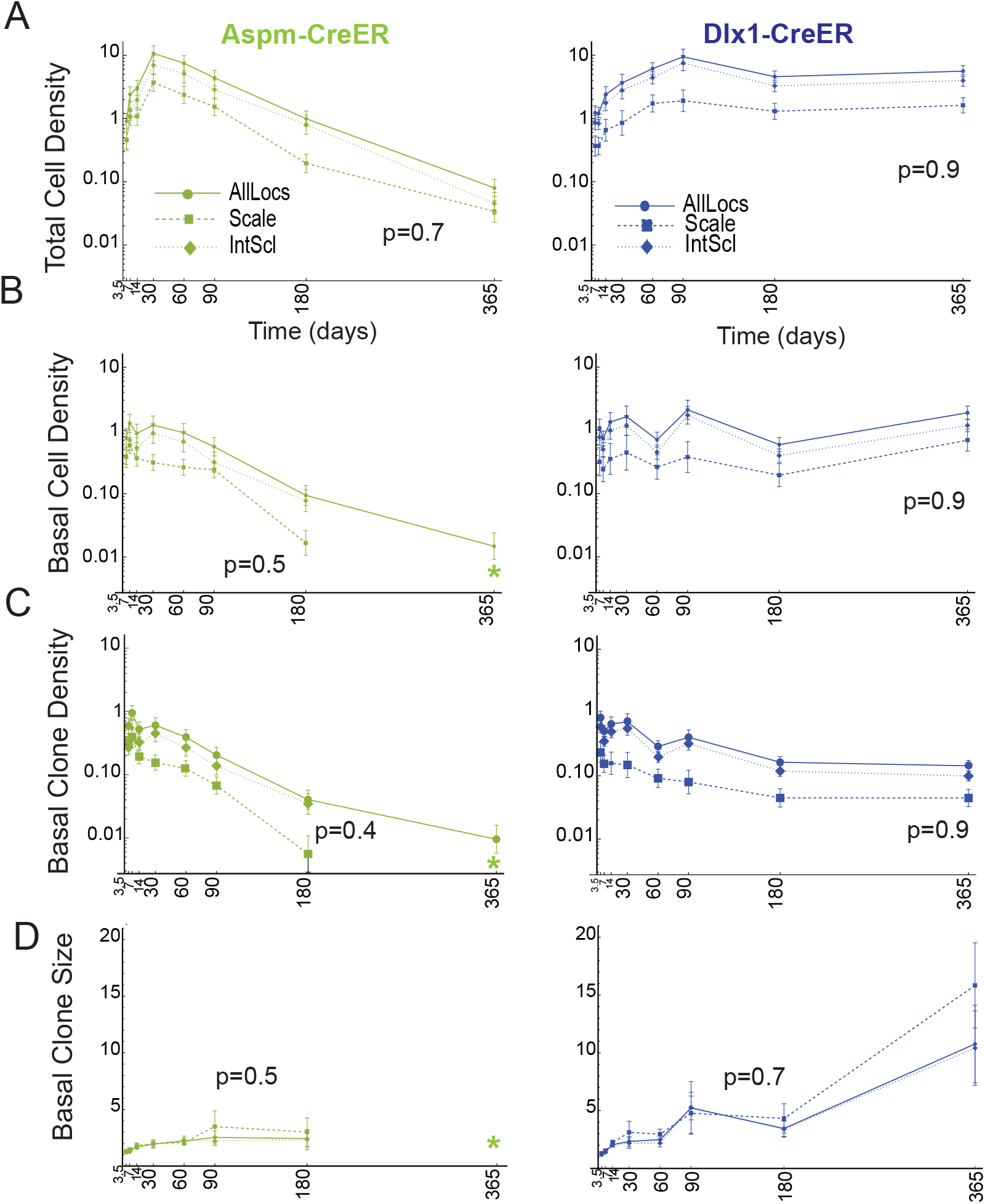
Two epidermal populations have homeostatic behaviors independent of location in scale vs inter-scale regions. Clonal data from Figure 2 is shown split in scale vs inter-scale regions of the mouse tail skin. No significant differences were observed between the two regions for each of the populations analyzed.

**Figure S6.**
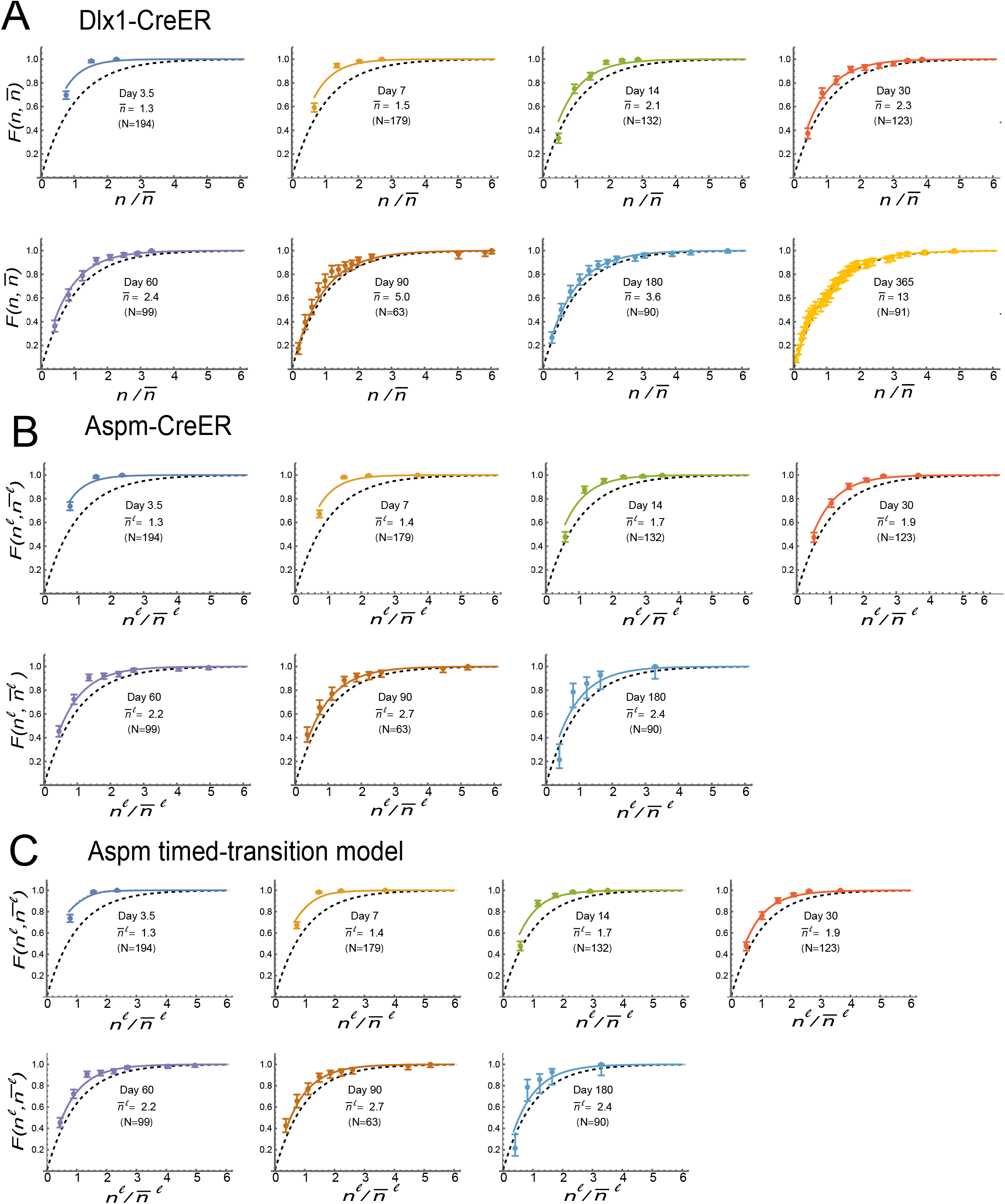
Neutral competition fits both Aspm-CreER and Dlx1-CreER population behavior. (A,B) The scaled clone-size cumulative plots for Dlx1-CreER and Aspm-CreER that are overlaid in Fig. 3D,E are displayed separately (see Fig 3 legend for more details). (C) The cumulative plots for the Aspm-CreER timed-transition model (Fig. 4). These are almost identical to the plots of the Aspm-CreER experimental data in (B).

## SUPPLEMENTARY MATERIALS AND METHODS

### Mouse care

All mouse work was executed according to Cornell University Institutional Animal Care and Use Committee guidelines (protocol number no. 2007-0125).

### Single cell RNA-seq data generation and processing for Pseudotime lineage trajectory analysis

Mice carrying pTRE-H2BGFP x K5tTA (*18*) were fed doxycycline supplemented chow as described and after 3 weeks of chase at PD52 single cell suspensions were isolated from back and tail skin as previously described (*19*). Three BL cellular subsets with distinct retention of H2B-GFP (~13,000 live cells total) were FACS sorted based on Sca1^+^/a6-integrin^+^ expression. The cells were subjected to sc-RNA sequencing according to the manufacturer’s protocol (10x Genomics). Approximately 270 M reads were generated for each of the 3 subsets from two replicates for both tail and back skin. High-quality cells that expressed about 200-5000 genes with a mitochondrial gene percentage under 10% were filtered. Scaling and expression normalization, PCA and UMAP dimensionality reductions and clustering were performed using the Seurat (*28*) R package (version 3.0.1). The samples were integrated using Harmony (version 1.0) (*29*) and nearly 13-14K BL cells were obtained after combining the three libraries. This combined dataset was subsequently used for all analysis in this study and this dataset is currently being deposited in GEO to make it accessible to public use (in progress). Trajectory analysis was performed on total cells by applying Monocle2 and *3*(*30*) to the Seurat (v3) output for the top 2,000 differentially expressed genes so as to assign Pseudotime values to individual cells. Cells that contained <200 transcripts were removed from further analysis. The trajectories were constructed with DDRTree. Feature plots were used in Seurat and Monocle2 and Monocle 3 to show Aspm/Mki67 expression in both mouse and human skin.

### Lineage tracing in whole mount tail epidermis

For lineage tracing, Dlx1-CreER (C57BL6)(*31*) (The Jackson Laboratory, no. 014551) or Aspm-CreER(*22, 32*), were crossed with Rosa–tdTomato reporter mice(*32*) (The Jackson Laboratory) and were genotyped as recommended by the manufacturer’s primer and protocol.

The clonal lineage tracing dose was one single injection of TM-100 μg/g body weight for Dlx1 and 10 μg/g body weight for Aspm-CreER mice at PD34. Mice were euthanized at the indicated times after the last injection. Non-injected CreER/Rosa–tdTomato mice were used to test CreER leakiness (not shown). For other experiments, Aspm-CreER were injected with 100 μg/g body weight TM for five consecutive days (high dose) or for 2 consecutive days (intermediate dose).

TdTomato^+^ cells were analyzed in whole-mount tail epidermis that was immunofluorescence (IF) stained to label the BL or to identify scale-interscale localization. To obtain this tissue, we separated the epidermis from the dermis; tail skin pieces (5 mm × 5 mm) were incubated in EDTA (20 mM)/PBS on a shaker at 37 °C for 2 h. Epidermis was then removed as an intact sheet followed by fixing in 4% paraformaldehyde (PFA) overnight at 4 °C. The skin pieces were washed, incubated in blocking buffer (1% BSA, 2.5% donkey serum, 2.5% goat serum, 0.8% Triton in PBS) for 3 h at room temperature, and incubated with primary antibodies/blocking buffer overnight at room temperature. Samples were washed 4× in PBS with 0.2% Tween for 1 h at room temperature, and were incubated overnight with secondary antibodies at 4 °C. After washing, samples were counterstained with Hoechst or DAPI for 1 hr and mounted.

#### Primary antibody dilutions

rat anti-β4-integrin (1:200, BD bioscience) or mouse anti-K10 (1:100, BioLegend no. 904301). All secondary antibodies (FITC, Cy5 or Alexa-594, Jackson ImmunoResearch) were used at a 1:500 dilution. Preparations were analyzed by confocal microscopy (Zeiss LSM710 or Zeiss LSM880) with Zen 2012 software. All confocal data are shown as projected *Z*-stack images viewed from the basal surface.

### Clonal analysis: counting of clones and cells within clones

Whole-mounts of tail epidermis obtained from lineage traced Dlx1-CreER or Aspm-CreER × Rosa-tdTomato mice induced at PD34 and stained for β4-integrin were imaged using confocal microscopy. Clones were analyzed at 3.5 days, 1 week, 2-week, 1 month, 2-months, 3-months, 6-months and 1-year post tamoxifen induction (Table S1). The number of tdTomato^+^ clones in the tail epidermis were counted on *Z*-stack confocal images (see data summary in Figure S1). Orthogonal views were used to display images in three dimensions to visualize the tdTomato positive cells and quantify the number of basal and total cells per clone. Cells were considered as basal when their basal side was positive for β4-integrin Each image was a stich of 30 tiles (e.g. xy fields of view with their corresponding Z-stacks). Clones were assigned in each tile and the basal and supra-basal cells were counted using the Zeiss Zenblue 2.5 software. Quantifications were independently performed at the various time points for each genotype (Table S1). More than 60 clones at each time-point were counted except for Aspm at 180 and 365 days, when clones were extremely rare or absent. Clone-size beeswarm plots (Fig. 2C,D were generated using R package 3.0.1.

In addition, for a macroscopic overview of clonal persistence (Fig. S4A,C), large areas of whole mount tail tissue (at least 6-8 images/mouse with at least 3 mice in each group) were scanned using the 10x objectives with a Leica Widefield fluorescence microscope, and the total number of clones present within the non-hair follicle area was counted.

### Adjustment for increasing tail area

Measurements of tail diameter and length from 14 to 365 days showed that there was a ~4% increase in tail area per month, independent of mouse gender. Cell and clone densities were adjusted for this small effect; the term “density” in the main text denotes “tail-area-adjusted density.”

### Data availability

The scRNA-seq raw data of both the replicates utilized in this paper have been recently reported (*20*) and deposited in the GEO database under accession code GEO: (*in process*).

---

**Table S1:** Summary of total and basal clone counts in the given number of mice and images quantified for the. Study:

| **Aspm-CreER** |  |  |  |  |  |  |  |  |
| --- | --- | --- | --- | --- | --- | --- | --- | --- |
|  | **3days** | **1 week** | **2 week** | **1month** | **2 months** | **3months** | **6months** | **1 year** |
| # Mice | 4 | 4 | 4 | 4 | 4 | 4 | 3 | 3 |
| # Images | 5 | 5 | 5 | 5 | 5 | 5 | 5 | 5 |
| # Tiles | 150 | 150 | **150** | 150 | 150 | 150 | 150 | 150 |
| Quantified by Sara | 0 | 0 | 0 | 0 | 0 | 0 | 0 | 0 |
| Quantified by Kevin | 150 | 150 | 150 | 150 | 150 | 150 | 150 | 150 |
| Total # Clones | 200 | 424 | 425 | 770 | 419 | 255 | 48 | 17 |
| Clones Basal | 167 | 256 | 150 | 174 | 123 | 66 | 15 | 3 |
| Clones Suprabasal | 33 | 168 | 275 | 596 | 296 | 189 | 33 | 14 |
| Clones Scale | 104 | 159 | 132 | 178 | 113 | 56 | 4 | 8 |
| Clones Interscale | 96 | 265 | 293 | 592 | 306 | 199 | 44 | 9 |

| **Dlx1-CreER** |  |  |  |  |  |  |  |
| --- | --- | --- | --- | --- | --- | --- | --- |
|  | **3days** | **1 week** | **2 week** | **1month** | **2 months** | **3months** | **1 year** |
| # Mice | 3 | 5 | 3 | 2 | 5 | 3 | 4 |
| # Images | 5 | 7 | 4 | 3 | 7 | 4 | 9 |
| # Tiles | 146 | 211 | **121** | 85 | 210 | 115 | 276 |
| Quantified by Sara | 116 | 151 | 91 | 55 | 120 | 85 | 186 |
| Quantified by Kevin | 30 | 60 | 30 | 30 | 90 | 30 | 90 |
| Total # Clones | 231 | 271 | 204 | 171 | 462 | 240 | 173 |
| Clones Basal | 218 | 192 | 153 | 132 | 118 | 88 | 107 |
| Clones Suprabasal | 13 | 79 | 51 | 39 | 344 | 152 | 66 |
| Clones Scale | 72 | 92 | 45 | 34 | 113 | 43 | 48 |
| Clones Interscale | 159 | 179 | 159 | 137 | 349 | 197 | 125 |

Basal clones are all clones that have at least 1 basal cell, and suprabasal clones are those with only suprabasal cells

**Table S2:** Rates and parameters calculated from lineage tracing data.

| Dlx1-CreER |  |  | Aspm-CreER |  |  |
| --- | --- | --- | --- | --- | --- |
| Definition | Parameter | Value | Value | Parameter | Definition |
| u-division ≈ u-export rate | $k$ ($\times 100/d$) | $2.3 \pm 0.5$ | $4.4 \pm 0.2$ | $k_+^{\ell}(0)$ ($\times 100/d$) | Observed amplification-phase u-division rate |
| Total division ≈ total export rate | $k^{\text{tot}}$ ($\times 100/d$) | $7 \pm 3$ | $21 \pm 7$ | $k_+^{\text{tot},\ell}(0)$ ($\times 100/d$) | Observed amplification-phase total division rate |
| Fraction of divisions that are coupled | $\frac{k^{\text{tot}} - k}{k^{\text{tot}}}$ | $0.7 \pm 0.1$ | $0.8 \pm 0.1$ | $\frac{k_+^{\text{tot},\ell}(0) - k_+^{\ell}(0)}{k_+^{\text{tot},\ell}(0)}$ | Observed fraction of coupled divisions in amplification-phase |
| u-rate imbalance (u-export–division) | $\Delta k$ ($\times 100/d$) | $-0.1 \pm 0.1$ | $1.4 \pm 0.2$ | $\Delta k(\text{ext})$ ($\times 100/d$) | Extinction-phase u-rate imbalance<br>$\Delta k(\text{ext}) = k_-(\text{ext}) - k_+(\text{ext})$ |
| | | | $3.5 \pm 0.8$ | $k_-(\text{ext})$ ($\times 100/d$) | Extinction-phase u-export rate |
| | | | $2.1 \pm 0.7$ | $k_+(\text{ext})$ ($\times 100/d$) | Extinction phase u-division rate |
| Half-life of population | $\tau_{1/2}$ | ∞ | $50 \pm 7$ | $\tau_{1/2}(\text{ext})$ (d) | Half-life of population in extinction phase |
| Maximum mean clone size | | ∞ | $2.5 \pm 0.5$ | $\bar{n}(\text{ext})$ | Maximum mean clone size in extinction-phase |
| Fraction of population replaced by nascent cells in homeostasis | | 0 | $\sim 0.01/d$ | | Fraction of population replaced by nascent cells in homeostasis |
| | | | $\sim 30\%$ | | Fraction of population in amplification-phase during homeostasis |

**Table S3.** Aspm-CreER ‘timed-transition’ model parameters and parameter predictions.

| Definition | Parameter | Value |
| --- | --- | --- |
| Nascent cell u-division rate | $k_+(0)$ ($\times 100/\text{day}$) | 4.5 |
| u-Division rate during extinction | $k_+(\text{ext})$ ($\times 100/\text{day}$) | 2.1 |
| u-export rate (constant) | $k_-$ ($\times 100/\text{day}$) | 3.5 |
| Transition time (amplification → extinction) | $\tau_{\text{transition}}$ (day) | 30 |
| Coupled:uncoupled division ratio (constant) | $k^c : k^+$ | 3.7 |
|  | **Prediction** |  |
| Fraction of the population input/day |  | 0.008 |
| Max mean # BL cells per nascent cell in amplification-phase | $\langle n \rangle_{\text{max}}$ | 1.3 |
| Mean exported cells per nascent cell in amplification-phase |  | 7.3 |
| Mean exported cells per nascent cell in extinction-phase (assuming constant coupled:uncoupled division rate ratio) |  | 12.7 |
| Fraction of cells in amplification-phase |  | 0.27 |
| Fraction of cells in extinction-phase |  | 0.73 |

## SUPPLEMENTARY THEORY

The use of Markovian birth-death processes to model the lineage tracing of self-renewing basal layer (BL) populations in homeostasis is well-established (for review, Refs. 14 and 15). Some division and BL→suprabasal layer (sBL) export events are correlated within clones on the timescale set by their rates (i.e., ‘coupled’, which could be due to asymmetric division [16] or correlated delamination and symmetric division [17]), but these leave the number of labeled BL cells unchanged and do not affect BL cell density, clone survival, or clone-size. Therefore, BL clone evolution depends only on the export-uncoupled (u)-division and division-uncoupled (u)-export rates, *k*_+_ and *k*_−_, respectively. (For notational simplicity, we group the possible event where both daughter cells from a division are immediately exported with the ‘uncoupled’ events.) We follow previous epidermal studies by modeling these processes as stochastic, cell-autonomous, and without microenvironmental influences, and test this assumption by examining the cumulative clone-size distributions [2–4, 14, 15]. sBL clonal evolution depends on potentially complicated transport process and, although rarely, sBL cells can migrate horizontally [17, 18], making associations with underlying BL clones difficult. Therefore, we only use the sBL lineage tracing data to estimate initial total division rates and instead focus our analysis on BL clonal evolution.

‘Critical’ birth-death models have often been used to analyze the lineage tracing of self-renewing (SR) populations in which the u-division and u-export rates are constant and equal (Ref. 14, for review). However, this type of model, which assumes balance between ‘birth’ (u-division) and ‘death’ (u-export), is not appropriate when the labeled subpopulation is non-self-renewing (NSR) with maturation-dependent growth rates, as observed here for Aspm-CreER. Non-self-renewal implies that the u-division and u-export processes are imbalanced, that the subpopulation is maintained by the constant input of new, *nascent* cells from a precursor, and that the growth properties of the nascent cells may change as they mature. In this case, the experimental lineage tracing observations will be averages that depend on the variation of labeling efficiency over the maturation-ages of the cells in the initial homeostatic population. This behavior was observed with Aspm-CreER. To analyze this data we adapted generalized non-Markovian birth-death modeling [19], which accommodates maturation-dependent rates, to lineage tracing with maturation-dependent labeling efficiency. Since this approach is new, we first present the theory in Sec. I, and then apply it to the experimental data in Sec. II. We assume that homeostasis is maintained throughout the chase period.

### I. GENERAL THEORY: HOMEOSTATIC NSR BL POPULATIONS IN LINEAGE TRACING

*A list of mathematical notations is in Sec. III.*

#### A. Labeled BL cell density of maturation-dependent NSR homeostatic populations in lineage tracing

Since the analysis of lineage tracing data from self-renewing populations using a critical birth-death model has been extensively discussed [14, 15], we focus on the analysis of data from NSR populations with maturation-dependent u-division and u-export rates and labeling efficiency—i.e., as is needed for analyzing Aspm-CreER data. We consider the critical model and the subcritical model for NSR populations with constant rates as limiting cases of the more general model. We define *maturity* τto be the time for which a cell lineage has evolved; i.e., since the introduction of its nascent cell founder.

##### 1. Behavior of a maturation-dependent NSR population in homeostasis

A NSR population requires a constant input of nascent cells for homeostasis and the properties of its descendants may change as the lineage matures. The mean number of descendants in the BL, 〈*n*〉, satisfies

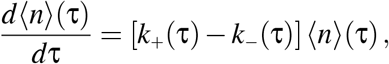

where *k*_+_(τ) and *k*_−_(τ) are the (finite) maturation-dependent u-division and u-export rates. Therefore,

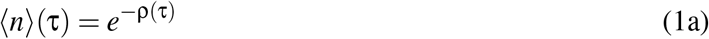

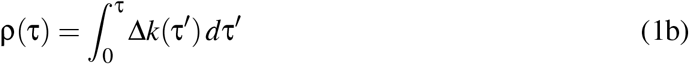

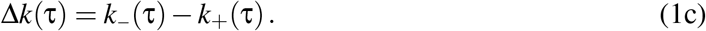

Δ*k*(τ) is the imbalance between the u-export and u-division rates at maturity τ.

In homeostasis, nascent cells are introduced into the population at a constant rate. Each begins its own lineage, so the steady-state population is a collection of lineages initiated at all times in the past; i.e., having maturities from 0 to ∞. Therefore, the probability that a cell in homeostasis has maturity τ is

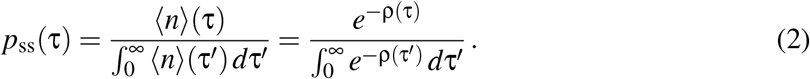

If cells are introduced at rate *r*_ss_ per unit area, the steady-state total BL density is

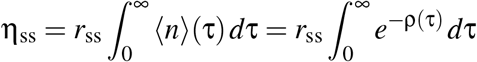

This density must be finite, which implies the necessary, though not sufficient, condition that

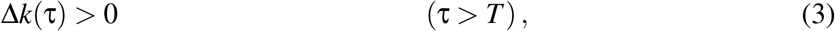

where *T* is a large value.

These equations, which describe the lineage of a nascent cell in homeostasis, set the stage for calculating the cell density and clonal evolution that will be observed for maturation-dependent NSR cells after labeling and lineage tracing.

##### 2. Observed BL density of NSR cells in lineage tracing

In a lineage tracing experiment, the steady-state population is labeled with relative efficiency ξ(τ), and this can affect the observations if it varies. To indicate this we attach the superscript^ℓ^ to all variables describing labeled cells.

The time evolution of η^ℓ^(τ, *t*), the density of labeled cells of maturity τ at chase-time *t*, satisfies

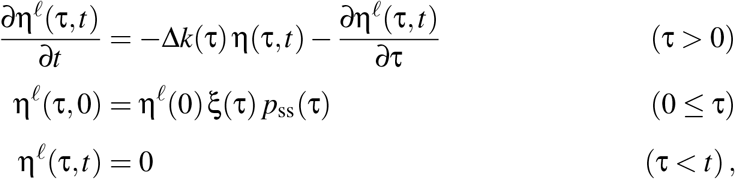

where, for convenience, we normalize *ξ*(τ) so that

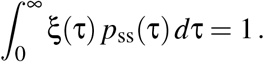

This has the solution

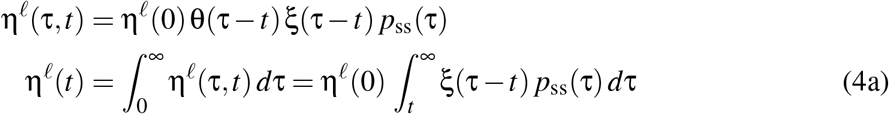

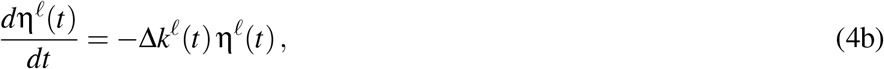

where η^ℓ^(*t*) is the total labeled cell density observed at chase-time *t* and

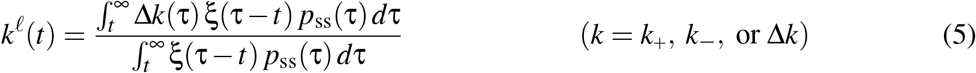

is the labeling efficiency-weighted (*lew*)-expectation of any rate at time *t*.

If *ξ*(τ) = *ξ* is constant,

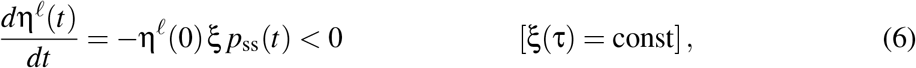

and η^ℓ^(*t*) will decrease monotonically with *t*, even when the ‘young’ cells are rapidly multiplying. This is because the loss of ‘old’ labeled cells, which must have Δ*k* > 0 (Eq. 3), will dominate any increase in the young labeled cells. However, *d*η^ℓ^(*t*)/*dt* can be positive at short times if young cells have Δ*k* < 0 and are preferentially labeled, as is the case with Aspm-CreER marked cells.

#### B. Generalized birth-death modeling of NSR population BL clone-sizes in lineage tracing

##### 1. BL clone-size distribution

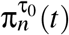, the probability that a cell of maturity τ_0_ at chase-time *t* = 0 evolves into a clone of size *n* ≥ 0 at time *t*, satisfies

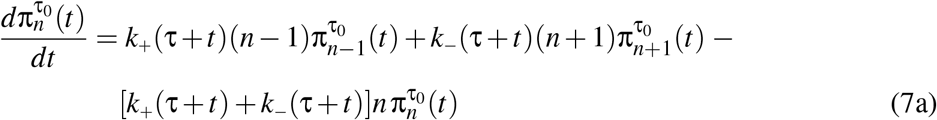

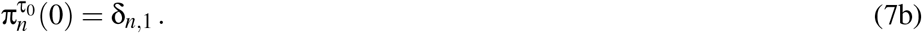

Kendall [19] solved these generalized birth-death equations for τ_0_ = 0. Extending his solution to τ_0_ ≥ 0 gives

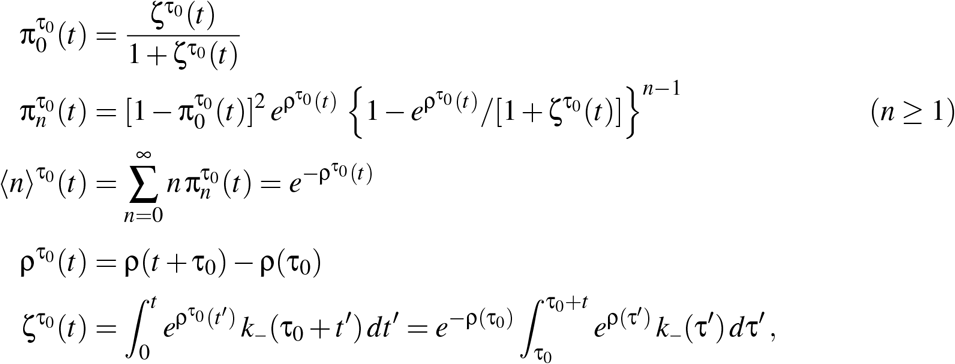

where 〈*n*〉τ_0_(*t*) is the mean clone-size including zero-size (i.e., non-surviving) clones of a lineage having maturity τ_0_ at *t* = 0. The probability that a clone with initial maturation τ_0_ survives (i.e., *n* 1) is

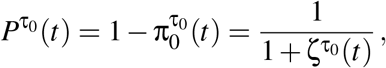

and the surviving clone-size probability distribution and mean surviving clone-size are

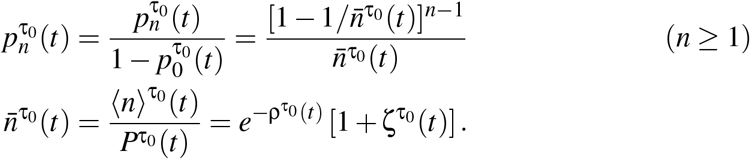

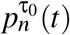 is a zero-modified geometric distribution which has the cumulative distribution

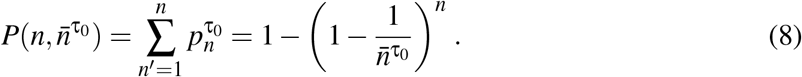

This simple distribution is the same for clones of any initial maturity τ_0_ at all chase-times, even when the u-division and u-export rates are maturation-dependent. It is a consequence of ‘neutral competition’—i.e., independent, random u-divisions and u-export of BL cells—and is predicted even in the absence of neutral drift. As we discuss below, while it is not the exact prediction for labeled cells in lineage tracing when the rates are maturation-dependent, it is a very good approximation when the spread in labeled τ_0_ is not large—i.e, the case with Aspm-CreER. When rates do not depend on maturity (e.g., Dlx1-CreER), Eq. 8 is exact and is equivalent to Eq. (1; main text).

When 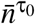 becomes large,

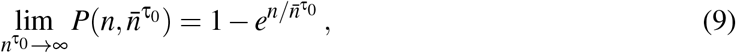

which is the ‘scaling limit’. It obtains as *t* → ∞ in the critical birth-death model for SC populations because the mean clone-size becomes arbitrarily large in this limit [14, 15]. However, in some cases the scaling limit will be observed without neutral drift. While the mean clone-size of a NSR population reach a finite maximum 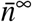 at large chase-times (see Sec. I C 2 below), the scaling limit will hold if 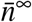 is sufficiently large. This is not the case for Aspm-CreER; therefore, its cumulative distribution never attains the scaling limit (Figs. 3, S6B, and S6C). However, in principle, 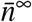 can be large even for a NSR population. Therefore, satisfaction of the scaling limit alone is a necessary but not sufficient condition for neutral drift.

##### 2. Observed BL clone-size distribution in lineage tracing

When labeling marks the steady-state distribution of cell maturities with efficiency *ξ*(τ), the probability distribution of labeled clone-sizes, including non-surviving, unobserved *n* = 0 clones, at chase-time *t* is 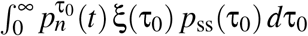. The probability that a labeled clone survives to be observed (i.e., *n* ≥ 1) is

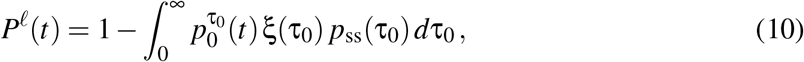

and the observed clone-size probability and mean clone-size are

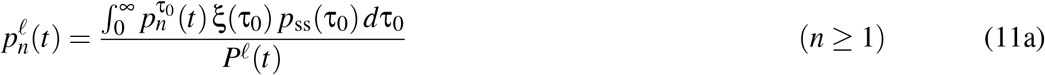

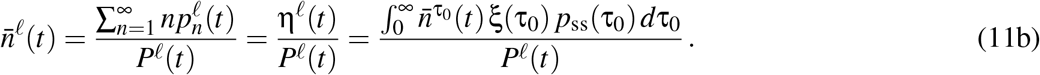

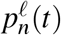 is a lew-average over zero-modified geometric distributions and is not exactly a zero-modified geometric distribution when the growth parameters are maturation-dependent. Therefore, Eq. (1; main text) will not be exact. However, it will be a good approximation at short times when 〈*n*〉^ℓ^(*t*) – 1 is small, because only 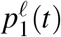 and 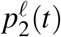 will be significantly bigger than zero, and these will be uniquely determined by 〈*n*〉 and 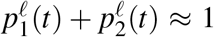. Eq. (1; main text) will also hold at long times if *k*_+_(τ) and *k*_−_ (τ) approach constants because the 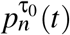 will become independent of τ_0_ in this regime (see Sec. I C 2). Eq. (1; main text) can also provide a good approximation at intermediate times, as is the case for the Aspm-CreER data and the timed-transition model discussed in Sec. II C (Figs. 3E and S6B,C).

#### C. Special cases

##### 1. Small t

Expanding the above equations in a Taylor’s series,

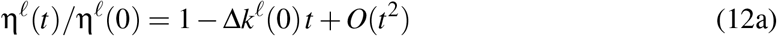

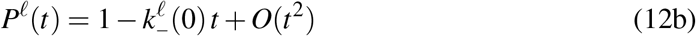

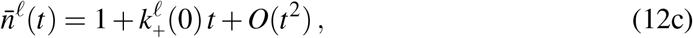

where 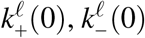, and Δ*k*^ℓ^(0), defined in Eq. 5, are the lew-expectations of the respective growth parameters over the homeostatic maturity distribution. We use Eq. 12c with 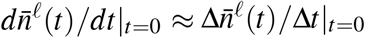 to determine 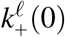 e.g., from the data in Figs. 3B,C. In principle, Eqs. 12a and b can be used to estimate Δ*k*^ℓ^(0) and 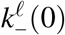, but we have found in practice that there is too much variability between microscopic images for accurate calculation of the required derivative.

##### 2. Large t

A case of special interest is when *k*_+_ (τ) and *k*_−_ (τ) asymptotically approach constants 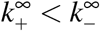 for *t* > *T*. This is true for Aspm-CreER in ‘extinction-phase.’ When *t* > *T*, all cells have τ_0_ +*t* > *T* (i.e., are ‘old’) and we can show that

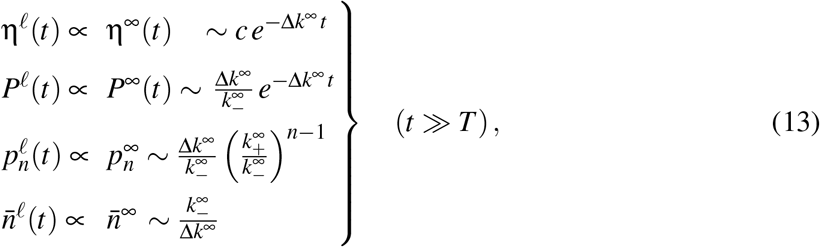

where *c* is a constant. The amount by which *t* must be greater than *T* for these approximations to hold depends in a complicated manner on *k*_+_(τ), *k*_−_ (τ), and *ξ*(τ); a conservative condition that is valid in all cases is Δ*k*∞(*t* – *T*) ≫ 1.

We use the superscript ^∞^ to indicate that the limits in this regime are independent of labeling efficiency because the rates are constant. We can use the extinction-phase measurements of *d*logη^ℓ^(*t*)/*dt* and 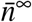 to compute Δ*k*∞, 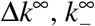, and (with Eq. 1c) 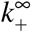. We call the values calculated for Aspm-CreER, Δ*k*(ext), *k*_−_(ext), and *k*_+_(ext).

In this situation, 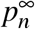 is a zero-modified geometric distribution. Therefore, Eqn. (1) in the main text is predicted to be exact for Aspm-CreER in extinction-phase.

##### 3. Constant k_+_ and k_−_; the subcritical case

The constancy of rates implies that, except for the overall normalization of η^ℓ^ the observations are independent of labeling efficiency and τ_0_. Δ*k* = Δ*k*^ℓ^ is a positive constant (see Eqs. 3 and 5), η^ℓ^(*t*) decreases exponentially (Eq. 4b), and evaluating Eqs. 10 and 11 gives^1^

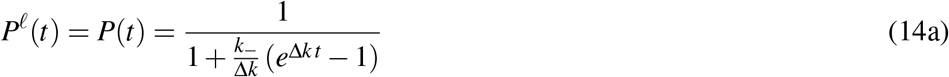

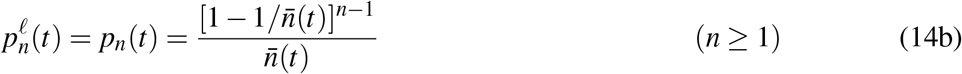

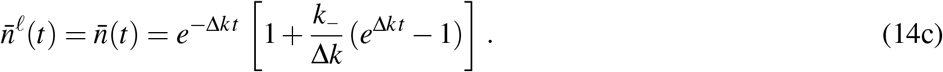

##### 4. Constant, equal rates: k_+_ = k_−_ = k; the critical case

Taking the limit Δ*k* → 0 in Eqs. 14 we get

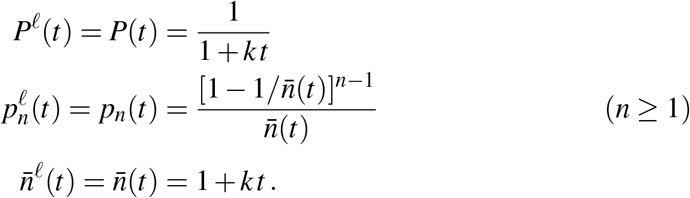

As in Sec. I C3, the observed rates are independent of labeling efficiency. This is the simple case of long-term SR cells (i.e., stem cells) where the critical birth-death model is applicable [14, 15]. The hyperbolic form of *P*(*t*) and the linear form of 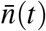 are used to fit the Dlx1-CreER clone survival and mean clone-size in Fig. 3B. The cumulative clone-size distribution is given by Eq. (1; main text).

### II. ANALYSIS OF THE EXPERIMENTAL RESULTS

#### A. Clonality

There was less than one labeled basal cell per 600 basal cells in all images at day 3.5. At this density, uniformly distributed cells would be well-spaced and 99% of the clusters should be clonal. Indeed, extrapolation of 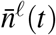 back to *t* = 0 for Aspm-CreER gave 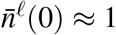, indicating that the marked cells were initially well-separated and that the subsequently observed clusters were clonal. However, the linear extrapolation for Dlx1-CreER gave 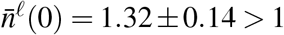 indicating (assuming that labeling was a Poisson process) that ~ 25% of the clusters were not clonal. This was probably because the Dlx1-CreER cells are spatially more restricted, being preferentially located in the interscale (Ref. 7 and Fig. S4), and the need to mark a sufficient number of cells for tracing made some non-clonality unavoidable. This does not affect η^ℓ^(*t*), but does affect 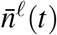, *P*^ℓ^(*t*), and 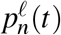 at the early time points. We accounted for this using appropriately modified boundary conditions when solving Eqs. 7 with *k*_+_ = *k*_−_.

#### B. Dlx1-CreER

The constancy of η^ℓ^(*t*) (Fig. 2E) implies that Dlx1-CreER marks a self-renewing population with Δ*k*^ℓ^ = (–0.001 ± 0.001)/d≈0. Consistent with the critical birth-death model and neutral drift, the density of labeled clones decreased hyperbolically (Fig. 3B), the mean labeled clone-size increased linearly (Fig. 3B), and the cumulative clone-size distributions agreed with Eq. (1; main text) (Fig. 3D and S6A). Thus, Dlx1-CreER appears to have constant, labeling-independent equal u-division and u-export rates, which makes it a SR stem cell population.

The linear best-fit to 〈*n*〉^ℓ^(*t*) and hyperbolic best-fit to *P*^ℓ^(*t*) (Fig. 3B) gave *k* = (0.023 ± 0.005)/d. We estimated the total division rate *k*^tot^ by comparing the early growth in total (BL plus sBL) cell density—i.e., before cells had time to be transported out of the sBL—with the BL density (Fig. S5A). Linear regression for the first 30 d gave 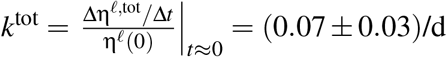, where the standard error estimate includes uncertaintis in the BL density. This suggests that (70 ± 10) % of the Dlx1-CreER divisions are coupled to export. However, the large experimental errors preclude exact quantitation.

#### C. Aspm-CreER

η^ℓ^(*t*) was approximately constant for the first 30 to 60 d and then began an exponential decrease during extinction-phase (Fig. 2F). This implies that the high-level Aspm-expressing cells that were labeled are nsr and that their growth properties depend on their maturity. We call the Aspm-CreER-marked cells ‘young’ or ‘old’ if their maturity is ≤ 30 d or ≥ 60 d, respectively, and draw the following conclusions:

##### a. Constant exponential rate of loss of old Aspm-CreER marked cells

*d*logη^ℓ^(*t*)/*dt* = –Δ*k*(ext) = – (0.014 ± 0.002)/d was constant during extinction-phase, and it is likely that 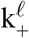 and 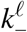 were constant as well. As discussed in Sec. I C 2, the constancy implies that the observed rates are independent of labeling efficiency and equal the actual rates. We used Eqs. 13 to compute 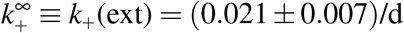 and 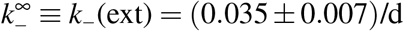 from Δ*k*(ext) and 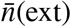.

##### b. Faster u-division in young cells

The biphasic behavior of the Aspm-CreER marked basal cell density cannot be explained as the mixture of two different labeled populations with maturation-independent growth parameters, since that would give a concave η^ℓ^(*t*) rather than the convex function observed. Using Eq. 12c, we calculated 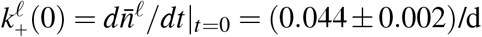. Because this is lew-averaged, the actual division rate of the young cells (i.e., with τ < 30 day) may be larger. Since this is twice *k*_+_(ext), it indicates that the u-division rate decreases with maturity.

##### c. Preferential labeling of young cells

If this were not so, Eq. 6 and the small value of *p*_ss_(*t*) = *d* η^ℓ^(*t*)/*dt* ≈ 0 when *t* ≤ 30 would imply that there were almost no young cells present in homeostasis, and the old cells would dominate the homeostatic labeled cell averages giving *k*_+_(0) ≈ *k*_+_(ext), which is false. We conclude that young cells with τ ≤ 30 d are preferentially labeled, possibly due to higher Aspm expression. In agreement with this, Aspm expression was observed to decrease in this lineage with increasing chase-time (Fig. S7).

##### d. Aspm-CreER young TA population expands its BL number

Because the slope of η^ℓ^(*t*) is small, the experimental error in the lineage tracing data precludes its accurate determination for *t* ≤ 30 d. η^ℓ^(*t*) is a time-shifted lew-average over 〈*n*〉(*τ*), the mean number of cells in the lineage of a nascent cell as it matures (Eqs. 1a, 2, and 4a). This ‘smears-out’ the change, so |*d*η^ℓ^(*t*)/*dt* | < |*d*〈*n*〉(τ)/*d*τ|_τ=*t*_| and the observed change in density is smaller than the change in the mean size of the nascent cell’s lineage. We conclude that 〈*n*〉(τ) increases for young cells during this early ‘amplification-phase’; i.e., Aspm-CreER is transit amplifying its basal cell numbers. Since the observed cell density underestimates the mean number of cells in a lineage, η^ℓ^(*t*) < 〈*n*〉 (τ) |_*τ*=*t*_, we conclude that, in homeostasis, at least 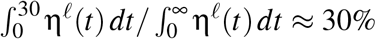 of the Aspm-CreER cells are in amplification-phase.

##### e. Aspm-CreER clonal evolution obeys neutral competition

The cumulative distributions 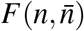 agree well with the neutral competition prediction, Eq. (1; main text) even though *ξ*(τ) varies. This is partly for the reasons discussed in Sec. I B 2. Another contributing factor may be that the preferential labeling of young cells means that only a relatively narrow maturation-window of cells, having a similar growth rates throughout the chase are labeled.

##### f. Most Aspm-CreER divisions are coupled to export in amplification phase

The initial labeling-averaged total division rate, calculated as described above using the *t* ≤ 30 d BL and sBL densities (Fig. S5B), was 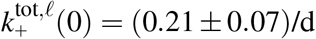. Since this is 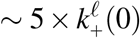, we conclude that most Aspm-CreER divisions are coupled to export.

##### g. More than ~ 17 cells may be exported for every nascent Aspm-CreER cell input

The rates calculated above and the behavior of η^ℓ^(*t*) implies that there are ~ 3 u-divisions over the lifetime of a nascent cell clone. In addition, ~ 6 coupled divisions occur during the first 30 days, so a minimum of 1 + 3 + 6 = 10 cells are exported. Furthermore, if the ratio 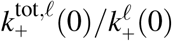 is constant during maturation, ~ 7 additional coupled divisions occur during extinction-phase so that ~ 17 cells are exported per nascent cell. This is an underestimate because it does not include the (unknown) amount of BL density increase in amplification phase and the reduction in the observed amplification phase division rates because of lew-averaging.

#### D. Aspm ‘timed-transition’ model

For this model we assume that Δ*k*(τ) changes from negative (cell density increasing) to positive (cell density decreasing) after some period of maturation because of a timed decrease in *k*_+_(τ) while *k*_−_ (τ) remains constant (Fig. 4A). For a simple approximation, we model the change as abrupt. To fit the data, we assume that crisis occurs at 30 days, that young cells with maturity τ ≤ 30 d have high 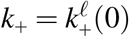 and enough Aspm to be labeled, while old cells with τ > 30 d have low *k*_+_ = *k*_+_(ext), lower Aspm expression, and are not labeled. Specifically,

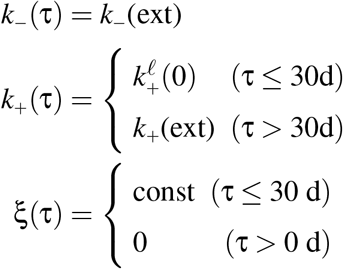

Applying the generalized birth-death model with these parameters gives a good fit to the data (Fig. 4B,C,D). Moreover, the scaled cumulative distributions predicted by the model agree closely with experiment (cf Figs. S6C), and both are very close to the predictions of Eqn. (1) in the main text. This model implies that about 1% of the Aspm-CreER lineage is replaced per day by the input of nascent cells and that 27% of the lineage is comprised of young cells. The mean number of descendants of a nascent cell in BL increases by ~ 30% before it begins to decrease at *t* = 30 d (i.e, 〈*n*〉_max_ = 1.3), but only a small ~ 5% increase in the labeled density η^ℓ^(*t*) is predicted by Eq. 4a, consistent with the experimental data, which is a lew-average. Each nascent cell undergoes about 3.5 u-divisions over the lifetime of its clone and, assuming that 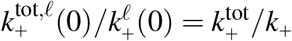 is constant, a total of ~ 20 cells are exported per input nascent cell.

### III. NOTATION

η^ℓ^(*t*): Labeled cell density at chase-time *t*
*k*_+_(τ), *k*_−_(τ): u-division, u-export rate of cell of maturity τ; u = uncoupled.
Δ*k*(τ): Rate imbalance for a cell of maturity τ; Δ*k*(τ) = *k*_−_ (τ) – *k*_+_(τ)
*k*^ℓ^(*t*): Lew-average of the corresponding division rate at chase-time *t*
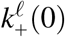: Observed amplification-phase lew-average of *k*_+_
*k*_+_ (ext), *k*_−_ (ext), Δ*k*(ext), 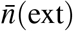: Observed (= true) values of these parameters in extinction-phase
lew: Labeling efficiency-weighted
〈*n*〉 (τ): Mean clone-size of maturity τ (includes zero-size clones)
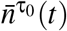: Mean surviving clone-size (*n* ≥ 1) at chase-time *t* of clones having initial maturity τ_0_
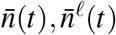: Mean surviving clone-size (*n* ≥ 1) at chase-time *t*
*p*_ss_(τ): Probability that a cell in homeostasis has maturity τ
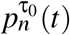: Probability (*n* ≥ 1) at chase-time *t* that a surviving clone of initial maturity τ_0_ has size *n*
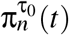: The probability (*n* ≥ 0) at chase-time *t* that a clone of initial maturity τ_0_ has size *n* (includes zero-size clones)
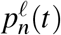: Observed lew-averaged probability (*n* ≥ 1) that a clone has size *n*
*P*^τ_0_^ (*t*): Probability that a clone of initial maturity τ_0_ survives to chase-time *t*
*P*(*t*), *P*^ℓ^(*t*): Clone survival probability at chase-time *t*; ^ℓ^ indicates that the observed value is a lew-average
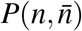: Cumulative clone-size, n, probability distribution for surviving clones of the same initial maturity having mean clone-size 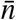
*r*_ss_: Rate of nascent cell density introduction during homoeostasis
ρ(τ): 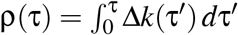
*t*: Chase-time
τ: Maturity
*ξ*(τ): Relative efficiency of labeling of cell having initial (*t* = 0) maturity τ

## SUPPLEMENTARY STATISTICAL METHODS

### I. CELL AND CLONE DENSITIES

The complete dataset for each population consisted of *N^a^* = 8 timepoint measurements in each of *N^x^* = 2 or 3 experiments. Each measurement contained cell counts from *N*^(*a,x*)^ images (typically 5 per experiment/timepoint), which were stitched together from 30 microscopic tiles as described in supplementary Materials and Methods. The datasets are described in Table S1. The variations between images were bigger than variations between mice, so images from all the mice in an experiment were pooled. Cell and clone counts varied between images because of both biological differences (e.g., location in the tail) and counting statistics. Therefore a compound distribution that accounts for both sources of variation was required; a standard Poisson-Gamma distribution was used. Another complication was that good spatial separation of clones required the frequency of tdTomato^+^ labeled cells to be very low, making it impractical to accurately determine the absolute labeling efficiency of each experiment. Therefore, we treated the absolute labeling efficiencies as statistical nuisance parameters that were removed using an empirical maximum marginal likelihood approach.

#### A. Poisson-Gamma distribution for counts in a single image

We modeled the biological variation of cell density between images using the Gamma distribution:

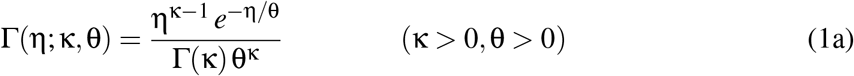

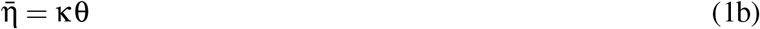

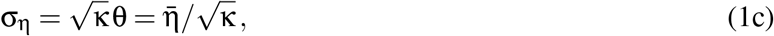

where 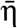 and σ are the expectation value and standard deviation of η, and κ and θ are the shape and scale parameters. The probability distribution of counts observed in an image of area *v* in an experiment with labeling efficiency *f* is the Poisson distribution

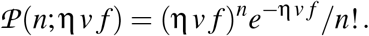

Convolving this with the Gamma distribution gives the Poisson-Gamma distribution:

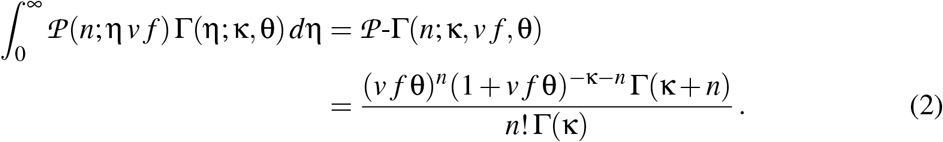

The variable of interest is the expected value of the density 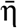, given by Eq. 1b. We expect the biological variations in density to be multiplicative, and so they are best analyzed on a log scale. Therefore, we reexpress the 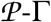 distribution as a function of *n, k, v f*, and 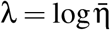 using Eq. 1b, and use a constant prior in **λ** for the maximum marginal likelihood analysis below.

#### B. The dataset likelihood

We denote the cell count for image *i* at timepoint *a* in experiment *x* as 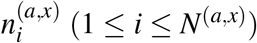, and the area of the image as 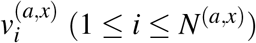. The goal is to determine the densities at the different timepoints, *λ^a^* (1 ≤ *a* ≤ *N^a^*). To avoid overfitting, we assume that the fractional biological variation in density, 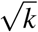 (see Eq. 1c), is similar for all measurements and fit the data with a single *k*. The log of the absolute labeling efficiency of experiment x was denoted as Δ^*x*^.

Denoting the set of data at timepoint *a* in experiment *x* as 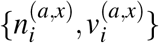, its log-likelihood is

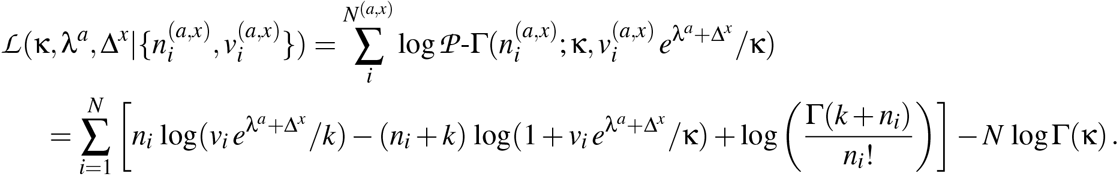

The log-likelihood for the complete dataset (i.e, from all timepoints in all experiments) is

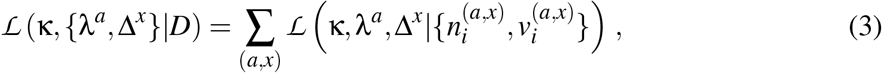

where *D* denotes the 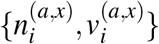 for all *a* and *x*.

Because the absolute labeling efficiency is unknown, only ratios between densities and labelling efficiencies—i.e., differences between the **λ**^*a*^ and Δ^*x*^ have significance, and 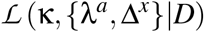 is degenerate under the combined transformation *λ^a^* → *c* + λ^*a*^, Δ^*x*^ → *c* – Δ^*x*^ (∀*a, x*), where *c* is an arbitrary constant. Without loss of generality, for computational convenience we temporarily break the degeneracy by fixing Δ^*Nx*^ = 0 and define the vectors

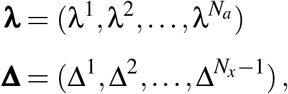

so that

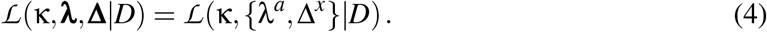

#### C. Empirical Bayes and maximum marginal likelihood

We first compute the maximum likelihood estimator (MLE)

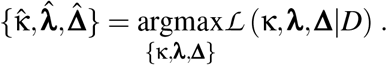

If the total number of measurements were much larger than *N^a^* + *N^x^* – 1 and all of the ∑_*x*_*N*^(*a,x*)^ were large, we could use the MLE and Hessian of 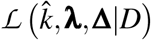 directly as estimators of the means and their covariance. However, when this is not the case, this approach may underestimate the statistical uncertainty in **λ**. A better estimate is obtained using the maximum marginal likelihood, For this purpose, we fix 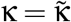, where

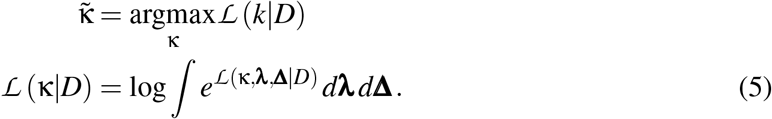

We have empirically found that each 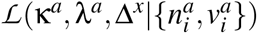 is close to quadratic in λ^*a*^ and Δ^*x*^ near its maximum, so it is appropriate to use a quadratic approximation to evaluate the complete integral in Eq. 5. 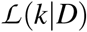 can then be numerically maximized to determine 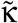. With **κ** fixed at 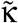, we then use a quadratic approximation to integrate over the nuisance vector Δ to get the marginal distribution of 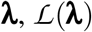, and compute its maximizer 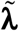.

#### D. Covariance matrix

As noted above, only relative differences between the 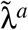 have significance; therefore, the marginal likelihood should be degenerate under under the transformation 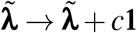, where **1** is the *N^a^*-vector with unity components and *c* is an arbitrary constant. This degeneracy was removed by fixing Δ*^Nx^* = 0 and is restored by projecting out the degenerate subspace, which is spanned by **1**.^2^ Therefore, the proper, degenerate covariance matrix is

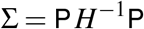

where *P* = *I* – **1** ⊗ **1** /*N^a^* is the projection operator with null-space **1**, and *H* is the *N^a^* × *N^a^* Hessian with elements

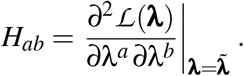

∑ was used in curve-fitting to the predictions of either the critical (Dlx1-CreER) or generalized (Aspm-CreER) birth-death models. This was performed using two-level random-effects models with restricted maximum likelihood estimation to account for both modeling and experimental errors [20]. The square-roots of the diagonal elements of ∑ were displayed as standard errors of the 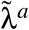 in plots; this is the standard error relative to the mean of the 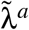.

#### E. Coordination of the experimental labeling efficiency estimates between different density calculations

The total cell density measurements provided the most accurate estimates of the variations in labeling efficiency between experiments. Therefore, we calculated a multinormal approximation to the marginal likelihood over Δ using the total cell density data, and then used the probability chain rule to calculate the likelihoods for the other densities in that population. Denoting the total cell density dataset as *D*^0^, the marginal likelihood over Δ computed with that dataset as *p*(Δ|*D*^0^), and the dataset being studied as *D*, we used

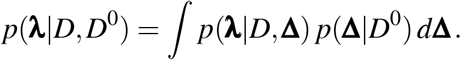

These integrals were evaluated using quadratic approximations as described above.

### II. CLONE SIZES

The variations in clone-sizes between images were larger than would be expected just from counting statistics indicating that there was additional biological variation. Therefore, mean clone-sizes were estimated using a two-level fixed plus random effects analysis [20]. Clone sizes were pooled for the scaled cumulative distribution plots.

Additional information on statistical methods is available upon request.

## Footnotes

1 In this simple case, the same results can be more directly obtained by solving Eqs. 7 using generating functions.

2 The same results would be obtained, albeit with more difficulty, if the computations had been performed using projections into the non-degenerate range of Q.

## REFERENCES

1. A. M. Klein, B. D. Simons, Universal patterns of stem cell fate in cycling adult tissues. Development 138, 3103–3111 (2011).

2. E. Clayton et al., A single type of progenitor cell maintains normal epidermis. Nature 446, 185–189 (2007).

3. X. Lim et al., Interfollicular epidermal stem cells self-renew via autocrine Wnt signaling. Science 342, 1226–1230 (2013).

4. G. Piedrafita et al., A single-progenitor model as the unifying paradigm of epidermal and esophageal epithelial maintenance in mice. Nat Commun 11, 1429 (2020).

5. P. Rompolas et al., Spatiotemporal coordination of stem cell commitment during epidermal homeostasis. Science, (2016).

6. K. R. Mesa et al., Homeostatic Epidermal Stem Cell Self-Renewal Is Driven by Local Differentiation. Cell Stem Cell 23, 677–686 e674 (2018).

7. G. Mascre et al., Distinct contribution of stem and progenitor cells to epidermal maintenance. Nature 489, 257–262 (2012).

8. A. Sanchez-Danes et al., Defining the clonal dynamics leading to mouse skin tumour initiation. Nature 536, 298–303 (2016).

9. M. Aragona et al., Mechanisms of stretch-mediated skin expansion at single-cell resolution. Nature 584, 268–273 (2020).

10. K. A. Cockburn et al., Gradual differentiation uncoupled from cell cycle exit generates heterogeneity in the epidermal stem cell layer. bioRxiv preprint doi: https://doi.org/10.1101/2021.01.07.425777, (2021).

11. L. Hayflick, P. S. Moorhead, The serial cultivation of human diploid cell strains. Exp Cell Res 25, 585–621 (1961).

12. E. Enzo et al., Single-keratinocyte transcriptomic analyses identify different clonal types and proliferative potential mediated by FOXM1 in human epidermal stem cells. Nat Commun 12, 2505 (2021).

13. C. S. Potten, The epidermal proliferative unit: the possible role of the central basal cell. Cell Tissue Kinet 7, 77–88 (1974).

14. C. S. Potten, M. Loeffler, Stem cells: attributes, cycles, spirals, pitfalls and uncertainties. Lessons for and from the crypt. Development 110, 1001–1020 (1990).

15. C. S. Potten, H. E. Wichmann, M. Loeffler, K. Dobek, D. Major, Evidence for discrete cell kinetic subpopulations in mouse epidermis based on mathematical analysis. Cell Tissue Kinet 15, 305–329 (1982).

16. D. G. Kendall, On the generalized “birth-and-death” process. Ann. Math. Statist. 19, 1–15 (1948).

17. C. Blanpain, Tracing the cellular origin of cancer. Nat Cell Biol 15, 126–134 (2013).

18. T. Tumbar et al., Defining the epithelial stem cell niche in skin. Science 303, 359–363 (2004).

19. A. Sada et al., Defining the cellular lineage hierarchy in the interfollicular epidermis of adult skin. Nat Cell Biol 18, 619–631 (2016).

20. S. Ghuwalewala et al., Epidermal basal domains organization highlights skin robustness to environmental exposure. bioRxiv https://doi.org/10.1101/2022.02.23.481662 (2022).

21. G. Chovatiya, S. Ghuwalewala, L. D. Walter, B. D. Cosgrove, T. Tumbar, High-resolution single-cell transcriptomics reveals heterogeneity of self-renewing hair follicle stem cells. Experimental dermatology 30, 457–471 (2021).

22. C. Marinaro et al., In vivo fate analysis reveals the multipotent and self-renewal features of embryonic AspM expressing cells. PloS one 6, e19419 (2011).

23. T. Lechler, E. Fuchs, Asymmetric cell divisions promote stratification and differentiation of mammalian skin. Nature 437, 275–280 (2005).

24. C. Blanpain, B. D. Simons, Unravelling stem cell dynamics by lineage tracing. Nat Rev Mol Cell Biol 14, 489–502 (2013).

25. Z. Lin et al., Murine interfollicular epidermal differentiation is gradualistic with GRHL3 controlling progression from stem to transition cell states. Nat Commun 11, 5434 (2020).

26. C. S. Potten, L. Kovacs, E. Hamilton, Continuous labelling studies on mouse skin and intestine. Cell Tissue Kinet 7, 271–283 (1974).

27. J. Lee et al., Runx1 and p21 synergistically limit the extent of hair follicle stem cell quiescence in vivo. Proceedings of the National Academy of Sciences of the United States of America 110, 4634–4639 (2013).

28. A. Butler, P. Hoffman, P. Smibert, E. Papalexi, R. Satija, Integrating single-cell transcriptomic data across different conditions, technologies, and species. Nat Biotechnol 36, 411–420 (2018).

29. I. Korsunsky et al., Fast, sensitive and accurate integration of single-cell data with Harmony. Nat Methods 16, 1289–1296 (2019).

30. C. Trapnell et al., The dynamics and regulators of cell fate decisions are revealed by pseudotemporal ordering of single cells. Nat Biotechnol 32, 381–386 (2014).

31. H. Taniguchi et al., A resource of Cre driver lines for genetic targeting of GABAergic neurons in cerebral cortex. Neuron 71, 995–1013 (2011).

32. L. Madisen et al., A robust and high-throughput Cre reporting and characterization system for the whole mouse brain. Nat Neurosci 13, 133–140 (2010).

## REFERENCES

[1] Lim, X. et al. Interfollicular epidermal stem cells self-renew via autocrine wnt signaling. Science 342, 1226–1230 (2013).

[2] Clayton, E. et al. A single type of progenitor cell maintains normal epidermis. Nature 446, 185–189 (2007).

[3] Mascre, G. et al. Distinct contribution of stem and progenitor cells to epidermal maintenance. Nature 489, 257–262 (2012).

[4] Sanchez-Danes, A. et al. Defining the clonal dynamics leading to mouse skin tumour initiation. Nature 536, 298–315 (2016).

[5] Wang, S. et al. Single cell transcriptomics of human epidermis identifies basal stem cell transition states. Nat. Commun. 11, 4239 (2020).

[6] Tumbar, T. et al. Defining the epithelial stem cell niche in skin. Science 303, 359–363 (2004).

[7] Sada, A. et al. Defining the cellular lineage hierarchy in the interfollicular epidermis of adult skin. Nat. Cell Biol. 18, 619–631 (2016).

[8] Butler, A., Hoffman, P., Smibert, P., Papalexi, E. & Satija, R. Integrating single-cell transcriptomic data across different conditions, technologies, and species. Nat. Biotechnol. 36, 411–420 (2018).

[9] Korsunsky, I. et al. Fast, sensitive and accurate integration of single-cell data with harmony. Nat. Methods 16, 1289–1296 (2019).

[10] Trapnell, C. et al. The dynamics and regulators of cell fate decisions are revealed by pseudotemporal ordering of single cells. Nat. Biotechnol. 32, 381–386 (2014).

[11] Taniguchi, H. et al. A resource of cre driver lines for genetic targeting of gabaergic neurons in cerebral cortex. Neuron 71, 995–1013 (2011).

[12] Madisen, L. et al. A robust and high-throughput cre reporting and characterization system for the whole mouse brain. Nat. Neurosci. 13, 133–140 (2010).

[13] Marinaro, C. et al. In vivo fate analysis reveals the multipotent and self-renewal features of embryonic aspm expressing cells. PloS One 6, e19419–e19419 (2011).

[14] Klein, A. & Simons, B. Universal patterns of stem cell fate in cycling adult tissues. Development 138, 3103–3111 (2011).

[15] Blanpain, C. & Simons, B. Unravelling stem cell cynamics by lineage tracing. Nat. Rev. Mol. Cell Biol. 14, 489–502 (2013).

[16] Lechler, T. & Fuchs, E. Asymmetric cell divisions promote stratification and differential of mammalian skin. Nature 437, 275–280 (2005).

[17] Rompolas, P. et al. Spatiotemporal coordinal of strem cell commitment during epidermal homeostasis. Science 352, 1471–1474 (2016).

[18] Ro, S. & Rannala, B. Evidence from the stop-EGFP mouse supports a niche-sharing model of epidermal proliferative units. Exp. Dermatol. 14, 838–842 (2005).

[19] Kendall, D. On the generalized “birth-and-death” process. Ann. Math. Statist. 19, 1–15 (1948).

[20] Raudenbush, S. Chapter 16, analyzing effect sizes:random-effects models. In Cooper, H., Hedges, L. & Valentine, J. (eds.) The Handbook of Research Synthesis and Meta-analysis, 295–315 (Russell Sage Foundation, 2009).

